# Can we study 3D grid codes non-invasively in the human brain? Methodological considerations and fMRI findings

**DOI:** 10.1101/282327

**Authors:** Misun Kim, Eleanor A. Maguire

## Abstract

Recent human functional magnetic resonance imaging (fMRI) and animal electrophysiology studies suggest that grid cells in entorhinal cortex are an efficient neural mechanism for encoding knowledge about the world, not only for spatial location but also for more abstract cognitive information. The world, be it physical or abstract, is often high-dimensional, but grid cells have been mainly studied on a simple two-dimensional (2D) plane. Recent theoretical studies have proposed how grid cells encode three-dimensional (3D) physical space, but it is unknown whether grid codes can be examined non-invasively in humans. Here, we investigated whether it was feasible to test different 3D grid models using fMRI based on the direction-modulated property of grid signals. In doing so, we developed interactive software to help researchers visualize 3D grid fields and predict grid activity in 3D as a function of movement directions. We found that a direction-modulated grid analysis was sensitive to one type of 3D grid model – a face-centred cubic (FCC) lattice model. As a proof of concept, we searched for 3D grid-like signals in human entorhinal cortex using a novel 3D virtual reality paradigm and a new fMRI analysis method. We found that signals in the left entorhinal cortex were explained by the FCC model. This is preliminary evidence for 3D grid codes in the human brain, notwithstanding the inherent methodological limitations of fMRI. We believe that our findings and software serve as a useful initial stepping-stone for studying grid cells in realistic 3D worlds and also, potentially, for interrogating abstract high-dimensional cognitive processes.

**Highlights:** We present software and an analysis method to probe 3D grid codes using human fMRI Based on an alignment score between 3D movement direction and grid orientation We then tested this using a 3D virtual environment and fMRI

Signals in entorhinal cortex were explained by a face-centred cubic lattice model

## Introduction

Grid cells in entorhinal cortex (EC) have received much attention from researchers in the field of spatial navigation because of their unique firing pattern. A grid cell, which is typically recorded in rodents when the animal explores a flat 2D surface in the laboratory, fires at multiple periodic locations resembling a hexagon (Hafting et al., 2005; Fig. 1A). Importantly, different grid cells have different spatial scales and phases, the combination of which enables efficient encoding of an entire space using relatively few cells, compared to when each individual cell fires at unique locations, as is the case with hippocampal place cells. Given that some animals, like bats, naturally explore volumetric space and humans can also explore 3D environments underwater or in microgravity conditions, the question naturally arises as to how grid cells would behave in 3D.

**Fig 1.**
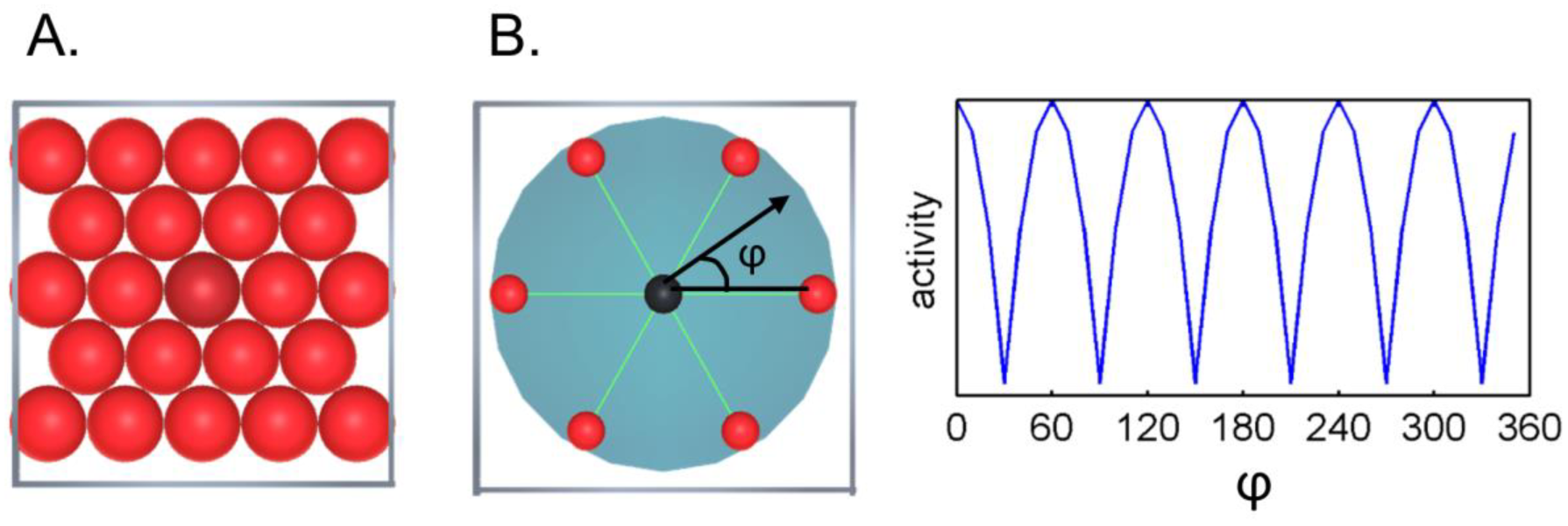
2D grid cells. **A.** A grid cell fires at multiple locations (called grid fields, red circles) which correspond to the centre of circles closely fitted in a 2D box. **B.** A grid cell’s activity is modulated by the animal’s movement direction (black arrow) relative to the grid axis (green lines linking one grid field to its neighbouring six grid fields). ϕ denotes the angle between the movement direction and the grid axis. The grid cell fires more when an animal’s moving direction is aligned to one of the grid axes. A grid axis is regularly displaced with 60° periodicity (right panel), therefore, a grid cell’s activity also shows a periodic response pattern depending on the animal’s movement direction.

The need to understand high dimensional grid codes has become more acute as a result of recent findings of grid cell involvement in non-spatial tasks. For example, electrophysiology studies have found that grid cells do not only encode the physical location of animals but also encode a continuously changing auditory tone in rats (Aronov et al. 2017) and visual space in primates (Killian et al. 2012). Human fMRI studies have also observed grid-like signals that encode locations during mental imagery (Bellmund et al., 2016; Horner et al., 2016), features of abstract visual stimuli (Constantinescu et al., 2016) and eye position during 2D visual search (Nau et al., 2018; Julian et al., 2018). This suggests that grid cells may be suitable for more abstract “cognitive mapping” (Tolman, 1948). If grid cells are indeed involved in abstract cognitive mapping, the ‘space’ might not be limited to simple 2D physical space on which most grid cell research has to date been conducted, because cognitive tasks can involve more than two features or attributes. Grid cells should also be able to efficiently encode 3D and higher dimensional space (unless the high dimensional cognitive problem can be projected into low dimensional space, e.g. context-dependent encoding).

Recent theoretical studies have offered predictions about the forms of grid codes that optimise encoding efficiency in 3D. These are analogous to the position of the centre of spheres tightly packed in 3D space, known as a face-centred cubic lattice (FCC), hexagonal close packing (HCP), or intermediate arrangements that yield the highest packing ratio (Mathis et al., 2015; Fig. 2). Although at least one research group is currently testing grid cells in flying bats (Ginosar et al., 2018), there is as yet no clear empirical evidence of grid cells showing a regular 3D structure. Technical difficulties associated with recording animals freely moving in 3D space might be one reason for the dearth of empirical findings relating to 3D grid fields.

**Fig. 2.**
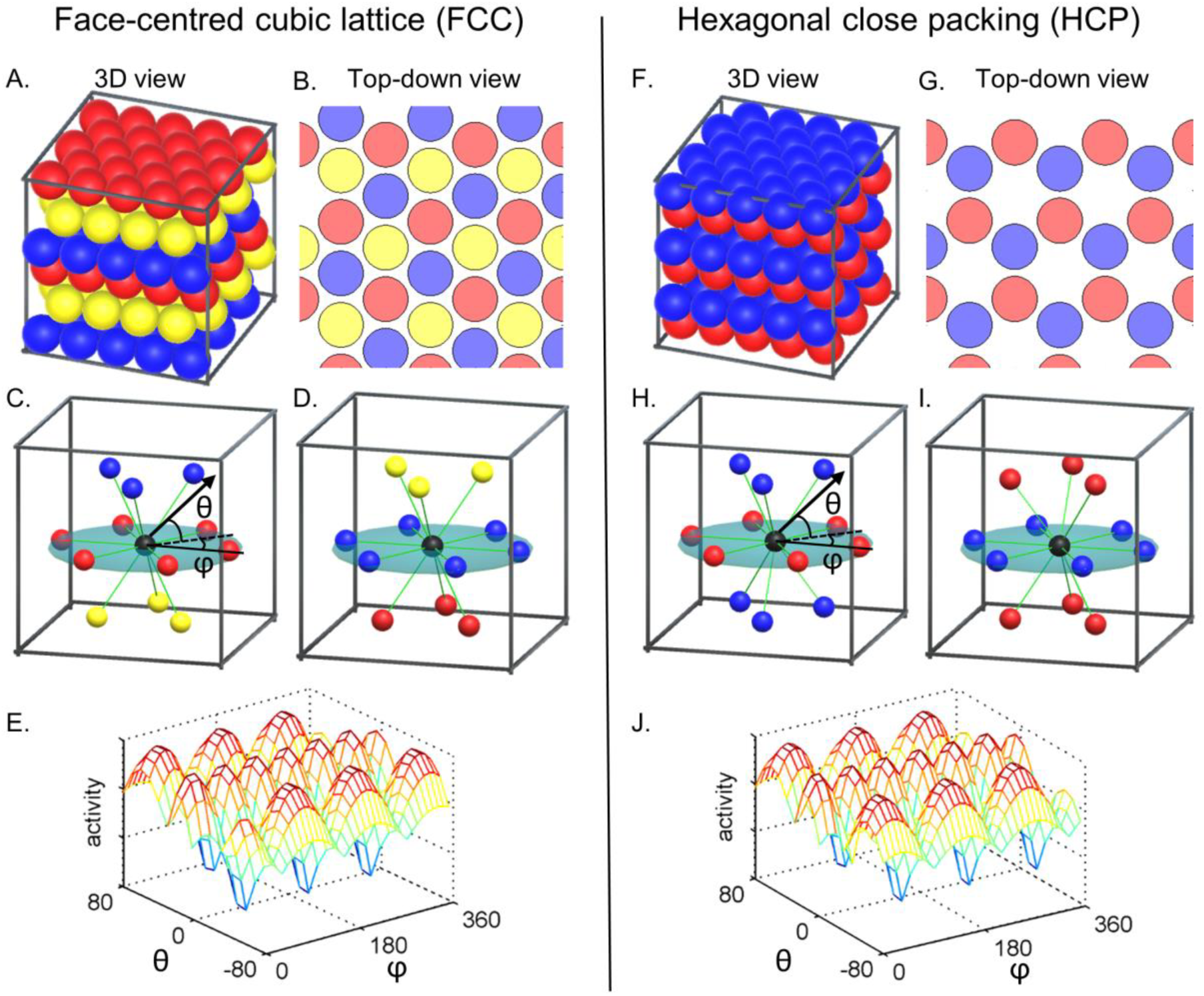
FCC and HCP models. A grid cell is proposed to fire at multiple 3D locations (called grid fields, coloured spheres) which correspond to the centre of spheres closely fitted in a box (**A,F**). The FCC and HCP arrangements have equally high packing density. The 3D arrangements can be viewed as 2D hexagonal lattices stacked on top of each other with a translational shift between the layers. FCC is composed of three repeating layers (blue, yellow and red spheres) and HCP is composed of two repeating layers (blue and red). The top-down views show spheres of reduced radius for visualization purposes (**B,G**). Grid axes are shown as green lines linking one grid field to the neighbouring 12 grid fields (**C,D,H,I**). FCC is a pure lattice in the mathematical sense, that is symmetric along the origin (e.g. blue and yellow spheres are symmetric; **C**), and can be described by three basis vectors. Thus, every sphere is surrounded by other spheres in the same arrangement, e.g. a sphere on the red layer (**C**) and a sphere on the blue layer (**D**). HCP lacks such symmetry (e.g. blue spheres on the top layer and bottom layer are not symmetric across the origin; **H**). The 3D arrangement of a unit cell is different for different layers, e.g. a sphere on the red layer (**H**) and a sphere on the blue layer (**I**) are surrounded by spheres that are 60° rotated from each other. Therefore, a grid axis can be defined only locally in HCP. Similar to a grid cell in 2D, it is expected that a grid cell’s activity is modulated by the alignment between an animal’s movement direction (black arrow) and the grid axis (green lines) (**C,D,H,I**). Depending on the vertical (θ) and horizontal (ϕ) component of the movement direction relative to the grid axis, a grid cell’s activity is expected to show a complex pattern of responses (**E,J**).

In humans, direct recording of grid cells in 3D is even more challenging. Invasive electrophysiology is only possible in a clinical setting with numerous limitations in experimental design, location of electrodes and the health of the participants. It would, therefore, be preferable to test for the existence of 3D grid codes non-invasively. Initial investigation with accessible and non-invasive methods like fMRI might also promote further research – for instance, the first intracranial recording of 2D grid cells in the human brain (Jacobs et al., 2013) was preceded by an fMRI study that found grid-like signals (Doeller et al., 2010). An obvious limitation of fMRI is that it cannot directly measure potential 3D grid fields. Yet, none of the previous theoretical studies on 3D grid cells suggested a method for detecting 3D grid cells using a macroscopic measurement technique like fMRI (Mathis et al., 2015; Stella and Treves, 2015; Horiuchi and Moss, 2015).

Therefore, the goal of this study was to examine whether it was feasible to empirically test different 3D grid models (e.g. FCC, HCP, or others) by extending the fMRI grid analysis that was originally developed in 2D (Doeller et al., 2010). In the Methods section, we first explain the principle of detecting 3D grid codes using the graphical user interface (GUI) software that we developed to visualize 3D grid structure and direction-modulated signals. A detailed description of the analysis method and several methodological issues that are either unique to 3D or relevant to both 2D and 3D then follows. Next we provide a proof of concept that our 3D grid analysis method can detect an FCC grid-like signal in the human EC by applying our analysis to empirical fMRI data. In this experiment, participants were moved in a virtual zero gravity environment inside the MRI scanner, with pre-scan training that involved a virtual reality (VR) head-mounted display. Finally, we discuss the methodological limitations and future directions for studying 3D grid signals.

## Methods

### Grid analysis

#### A principle for detecting 3D grid codes using fMRI

Before we propose a method for detecting 3D grid codes, we will first summarise how 2D grid codes have been probed in previous fMRI studies. fMRI measures the gross activity of thousands of neurons via complex neural-hemodynamic coupling. When the thousands of grid cells that fire at different locations are summed up, the gross activity is no longer expected to respond to fixed periodic locations in the environment. However, there is another important property of grid cells that enables their detection at a macroscopic level like fMRI (Doeller et al., 2010). The activity of ‘conjunctive’ grid cells is modulated by the alignment between the movement direction of an animal and the grid axis (Doeller et al., 2010). This means that a grid cell shows greater activity when an animal moves in directions parallel to one of the grid axes compared to other directions (Fig. 1B). As the majority of grid cells share a common grid axis, the summed response of thousands of grid cells can be systematically modulated by the movement direction of a participant. In previous studies that investigated grid cells in 2D, fMRI activity was modelled as a cosine function of movement direction relative to the grid axis with a period of 60 degrees to account for hexagonal symmetry (e.g. Doeller et al., 2010; Constantinescu et al., 2016). Another explanation for the direction-modulated grid signal measured by fMRI is that when a participant’s movement direction is aligned with the main grid axis, relatively few grid cells are repeatedly activated, whereas when the movement is not aligned with the main grid axis, more cells are irregularly activated. The fMRI response can be different in these two cases due to non-linear neural-hemodynamic coupling (e.g. Doeller et al., 2010).

We assume that the same principle of direction-modulation will hold in 3D, so that fMRI activity can be modelled as the degree of alignment between 3D movement direction and the grid axis. The question we face is how to define and predict the grid axis if grid cells have 3D receptive fields following either the FCC or HCP arrangements (Mathis et al., 2015; Stella and Treves, 2015). Both FCC and HCP arrangements are analogous to the spatial arrangement of tightly stacked spheres inside a box with a minimum gap (e.g. like oranges in a crate). These 3D arrangements can be viewed as 2D hexagonal lattices stacked on top of each other with a translational shift between the layers. FCC is composed of three repeating layers (blue, yellow and red spheres, see Fig. 2A,B) and HCP is composed of two repeating layers (blue and red spheres, see Fig. 2F,G). In both models, one centre sphere is surrounded by 12 neighbouring spheres (Fig. 2C,D,H,I). To enable ourselves and other researchers to visualize the 3D receptive fields of grid cells, we developed interactive web-based software where users can zoom, pan, rotate, and cross-sect the 3D arrangements (the software, including a manual, can be accessed here: www.fil.ion.ucl.ac.uk/Maguire/grid3D_gui). Example screenshots are shown in Fig. 3. The software was implemented using Unity 5.4 (Unity Technologies, CA, United States). Of note, this software was developed for visualization purposes and does not have an fMRI data analysis function. fMRI analysis software for 2D grid codes is already available (Stangl et al., 2017).

**Fig. 3.**
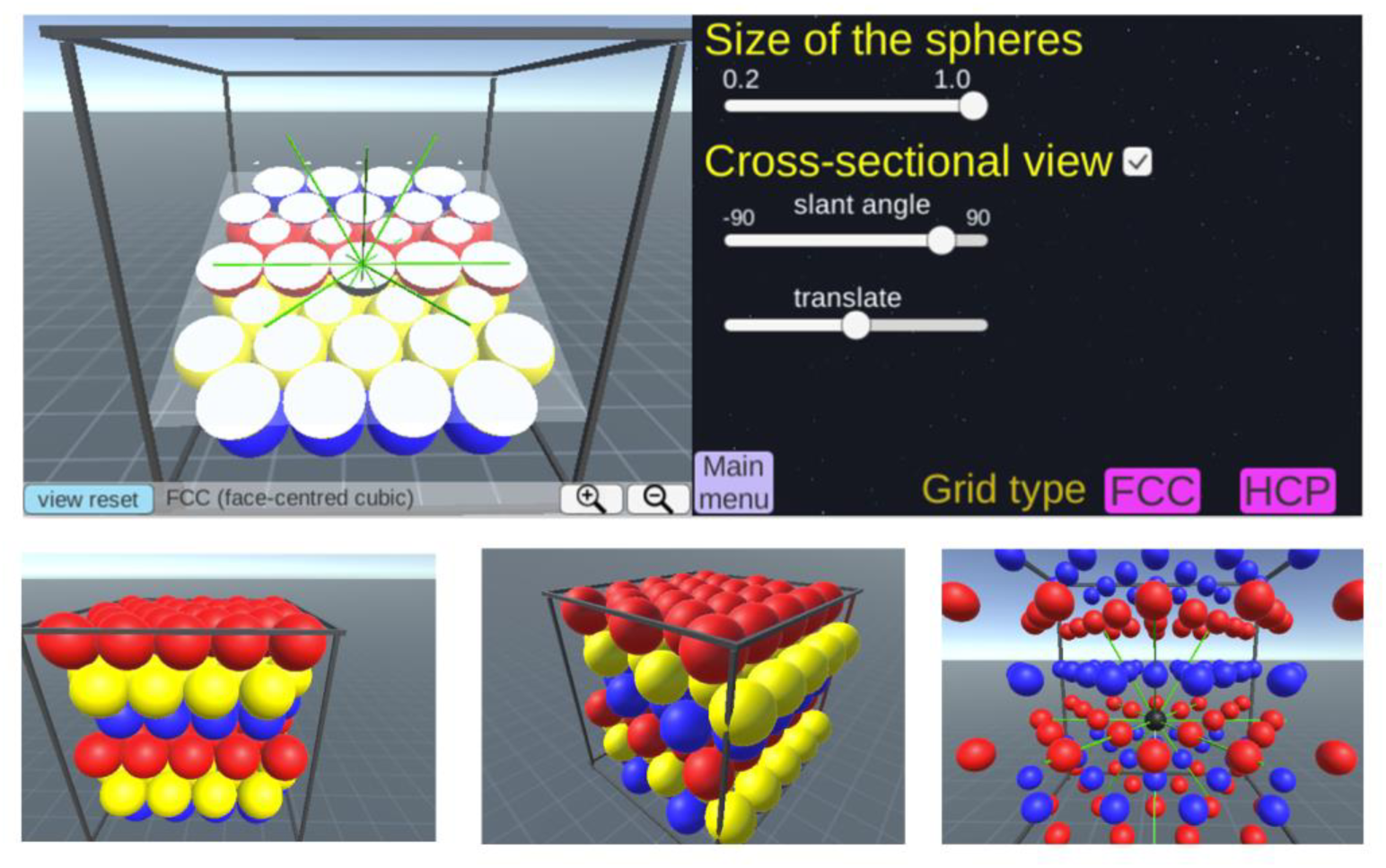
Screenshots of our 3D grid fields visualization software. The software, including a manual, can be accessed here: www.fil.ion.ucl.ac.uk/Maguire/grid3D_gui. Users can change the viewpoint, the size of the spheres, and select cross-sectional views of grid fields. Users can also switch between the FCC and HCP arrangements.

FCC is a pure lattice where the location of each node can be described by a linear combination of three basis vectors. The FCC arrangement is symmetric across the origin (e.g. the blue and yellow spheres are facing each other, Fig. 2C) and every grid field is surrounded by 12 grid fields in the same arrangement. Thus, it is intuitive to define the grid axis as the direction linking one grid field to its neighbouring grid fields (the green lines linking a centre black sphere to neighbouring coloured spheres in Fig. 2C,D). We can then model the fMRI signal as a cosine of the angle between the 3D movement direction and the nearest grid axis. Thus, a larger signal is expected when the angle is smaller (i.e. the movement is aligned to the grid axis). The nearest grid axis forms the minimum angle with the direction vector. We can predict the grid cell’s response when a participant is moving in a particular 3D direction, defined by azimuth (horizontal angle) and pitch (vertical angle) using our interactive software (example screenshots are shown in Fig. 4A,B). Figure 2E shows the predicted grid activity as a function of azimuth and pitch of a movement direction relative to the grid axis.

**Fig. 4.**
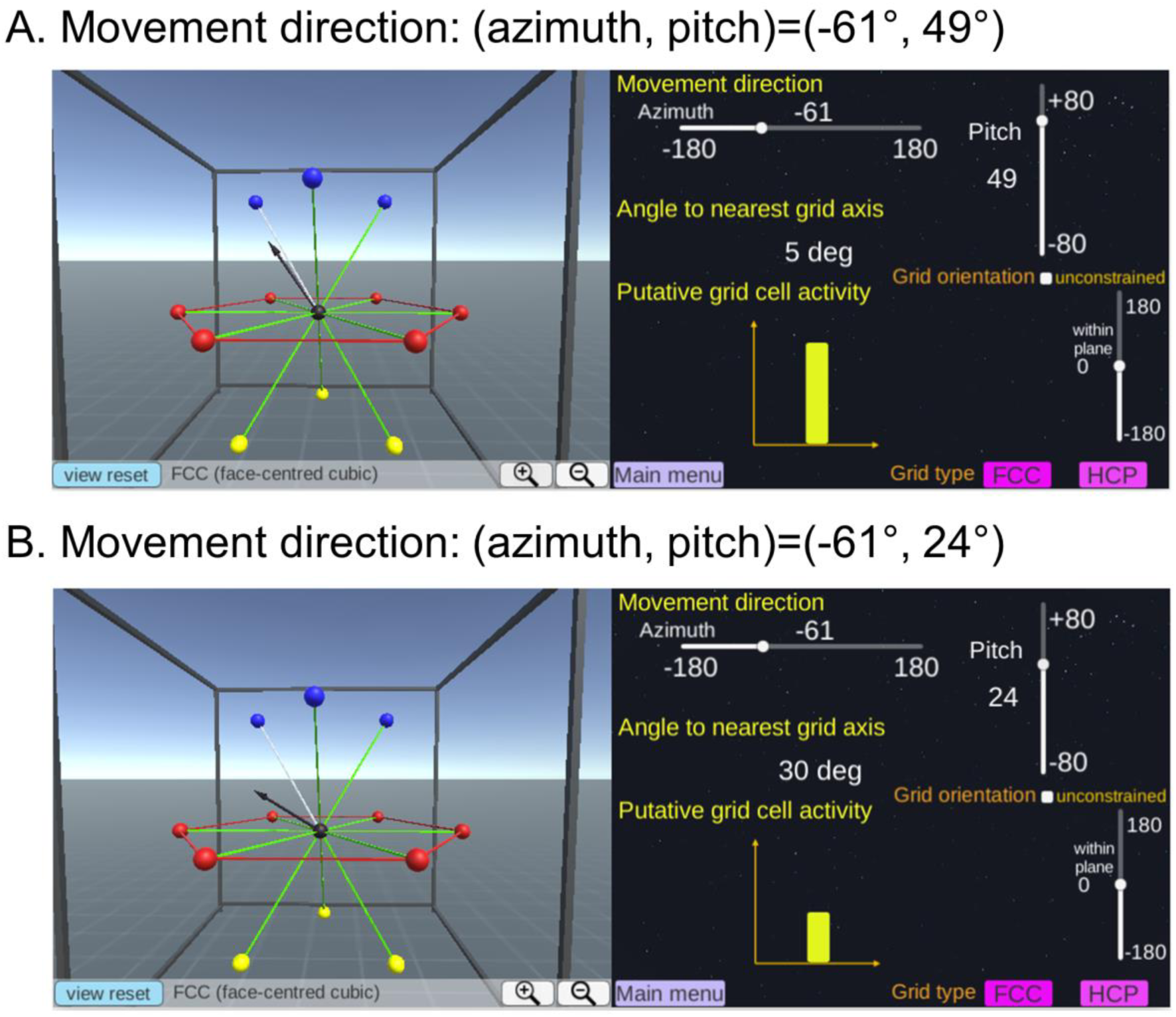
A grid cell’s activity is modulated by the movement direction relative to the grid axis. **A.** A participant’s 3D movement direction (black arrow, left panel) is close to the grid axis (white line, left panel) with 5° deviation. Thus, the activity of grid cells is expected to be high (yellow bar graph, right panel). **B.** A participant’s movement direction (black arrow, left panel) is far away from the grid axis (white line, left panel) with 30° deviation. Consequently, less activity is expected (yellow bar graph, right panel). Users can change the movement direction using the sliders on the right panel. Users can also switch between the FCC and HCP models and change the viewpoint.

Importantly, predicting the direction-modulated grid signal is not straightforward for the HCP model because HCP is not a pure lattice, as has been discussed in previous theoretical studies (Mathis et al., 2015; Stella and Treves, 2015). Unlike the FCC, the HCP arrangement is symmetric across the horizontal plane (e.g. the blue spheres on a layer above the red spheres are located at the identical positions as the blue spheres on the layer below, Fig. 2H). Therefore, the arrangement of unit grid fields is dependent upon the layer (Fig. 2H and Fig. 2I are 60° rotated from each other). This means that the grid axis is defined only locally in HCP (Supplementary Fig. 1; note that this is more evident when viewing the 3D arrangements using our 3D visualization software). This imposes a limitation in predicting a direction-modulated fMRI signal which is dependent upon the global grid axis. There is currently no empirical data on how the fMRI signal would vary when receptive fields of grid cells follow a non-lattice structure. One can, nevertheless, attempt to model the grid voxel’s activity for the HCP model using one set of locally defined grid axes (Fig. 2) with this caveat in mind. This yields largely similar response patterns to the FCC model except for the difference in vertical symmetry (Fig. 2E and J). We later show a simulation whereby the FCC and HCP models can still be distinguished if the HCP model follows the locally defined grid axis (see the section “Estimating 3D grid orientation”). Predicting fMRI signals is even more challenging if grid cells have 3D receptive fields that follow neither FCC nor HCP models. 2D hexagonal layers that are stacked with any order (e.g. blue – yellow – red – yellow – red – blue layers in Fig. 2A) also yield the same packing density and have been proposed as a probable grid code (Mathis et al., 2015). In this case, there is more variation in the local grid axis.

In summary, we predict that a macroscopic measurement which relies upon the shared grid axis of neighbouring grid cells would be most sensitive to detecting a grid response that follows a perfect lattice structure – FCC.This means that a macroscopic measurement that relies upon the direction-modulation of a grid signal is unfortunately not suited to making comparisons between different 3D grid models. Nevertheless, developing an analysis method that can probe at least one type of 3D grid model using fMRI is an important starting point for understanding whether 3D grid codes are even detectable in humans. In the rest of the Methods section, we detail the analysis method and validation with this purpose in mind.

#### The orientation of the grid axis relative to the environment

In the previous section, we explained that grid activity can be modelled as movement direction relative to the grid axis. Crucially, the movement direction of a participant is known to experimenters, while the orientation of the grid axis relative to the 3D environment is unknown. Fig. 5 describes two hypothetical cases where a participant moves in the same direction but the grid axis is oriented differently. In Fig. 5A, a participant’s movement direction (black arrow) is relatively closely aligned to the grid axis with an angular deviation of 14°, resulting in high activity (the yellow bar graph). In Fig. 5B, due to a different orientation of the grid axis, the same movement direction is farther away from the grid axis with an angle of 35°, resulting in low activity. The orientation of the grid axis can be numerically estimated by iteratively fitting the experimental data - a process we describe in a later section.

**Fig. 5.**
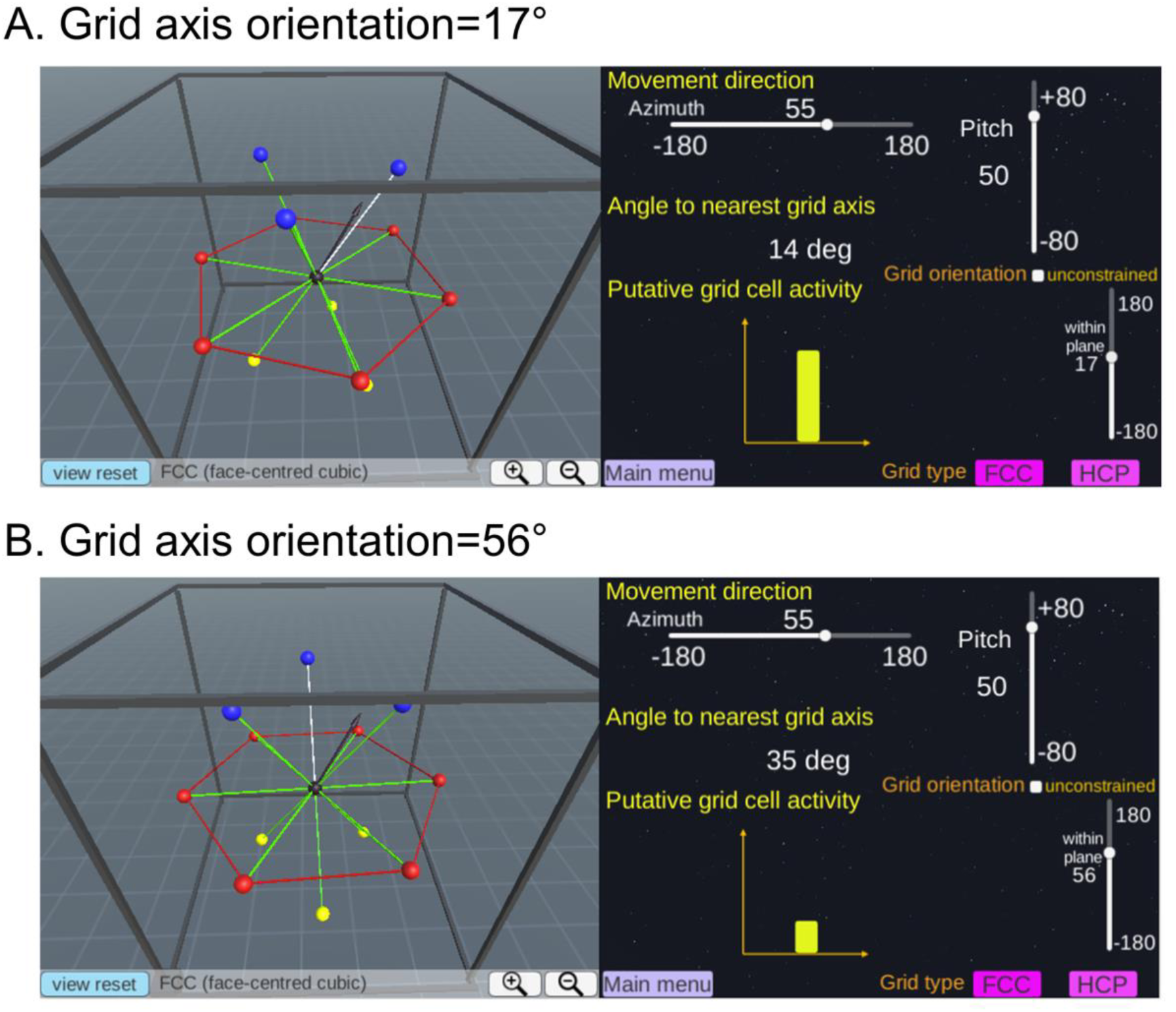
The orientation of the grid axis relative to the environment. The movement direction (black arrow, left panel) is identical in **A** and **B** (azimuth=55°, pitch=50° in this example). However, grid cells are aligned differently relative to the environment (rectangular frame, left panels in **A** and **B**), meaning that the grid axes are rotated from each other (left panel). Thus, the grid alignment scores measured as the angle between the movement direction and the nearest grid axis differ (14° versus 35°), resulting in different amounts of grid activity (yellow bars graph, right panels in **A** and **B**). Users can change the orientation of the grid axis using the sliders on the right panel.

Unlike in 2D where the orientation of the grid axis can be specified by one polar angle from a reference direction (e.g. 20° from the north-south axis), the grid axis in 3D can in theory be rotated along any three arbitrary axes and the order of applying each rotation also matters (known as the non-commutative property of 3D rotation). Although users can explore these 3D rotation options in our software, we restricted the rotation of the 3D grid axis to only one axis so that six hexagonal grid fields (the red spheres in Fig. 5) remain parallel to the ground of the environment when we analyse our fMRI data. This restriction in the rotation axis can be justified by the fact that grid cells on a 2D horizontal surface show corresponding grid fields. This restriction is also required when comparing putative grid orientations across multiple voxels or multiple participants using a standard circular statistic like a von Mises distribution. Conducting the modelling process in this way simplifies it and reduces the computational cost and the risk of overfitting GLMs for hundreds of possible 3D rotations with a limited fMRI time series.

#### The relationship between the grid alignment score and fMRI activity

A grid voxel’s activity is expected to be modulated by the degree of alignment between the movement direction and the grid axis, and we defined the grid alignment score as the cosine of the angle between the movement direction and the nearest grid axis. This is equivalent to the previous grid analysis in 2D which used parametric regressors of cosine and sine functions (Doeller et al., 2010; Horner et al., 2016), except for a minor difference in that our method predicts more abrupt changes in grid alignment scores when the movement direction passes the midpoint of two neighbouring grid axes (Supplementary Fig. 2B,C). Of note, the precise form of the direction-modulated firing rate of grid cells is not known in either 2D or 3D. There is also additional complexity in measuring a grid cell’s signal via neural-hemodynamic coupling. Therefore, it is also possible to model the grid voxel’s activity with a non-sinusoidal function, like a linear, binary or even more complex functions (Supplementary Fig. 2D,E). The exact relationship between direction-modulated grid activity and the fMRI response should be examined in future studies, but for now we believe that our choice of modelling the fMRI signal with the cosine of the angle is a reasonable starting point, as it is in line with previous studies in 2D.

#### Estimating 3D grid orientation in fMRI data

In this section, we describe how to estimate 3D grid orientation from an fMRI time series. As explained earlier, the grid alignment score can be calculated as the cosine of the angle between the participant’s movement direction (known to experimenters) and the nearest grid axis. The nearest grid axis is determined by the orientation of the grid axis relative to the environment, which is unknown to the experimenters (Fig. 5). In 2D, the grid activity can be modelled as a simple cosine function of the horizontal movement direction (ϕ) and the orientation of grid axis (ω), and the orientation (ω) can be estimated analytically by fitting cosine and sine functions in a GLM (e.g. a quadrature filter, Doeller et al., 2010):

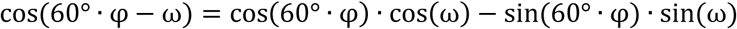

However, a simple analytical function of azimuth (ϕ), pitch (θ) and the grid axis orientation (ω or more parameters) which can describe the grid alignment score in 3D is unknown. We, therefore, suggest a simple numerical method to estimate grid orientation as follows:

We first assume that the grid orientation is aligned at 0 degrees from a reference direction (e.g. parallel to the side wall of the environment) and then calculate the grid alignment score. This grid alignment vector is then convolved with the hemodynamic response function (SPM canonical hemodynamic response function). The resulting vector serves as a hypothetical grid voxel signal. We create a general linear model (GLM) which contains this predictive 3D grid signal and nuisance regressors that include six head motion realignment parameters and experiment-specific conditions like an occasional question and response period. The fMRI time series (after standard preprocessing) in each voxel and in each scanning session is then fitted with the GLM, and the outcomes - beta (regression coefficients) and adjusted R square values (Soch et al., 2018) - are saved for each voxel.

We then repeat the whole procedure with newly calculated grid alignment scores with different assumptions, namely that the grid orientation is aligned at 15, 30, 45,…, 120 degrees relative to the environment (note that we only need to sample the grid orientation up to 120 degrees, as the geometry of the 3D lattice structure of both FCC and HCP is symmetric for the 120 degree rotations on a plane). For each voxel and each scanning session, we select the orientation of the grid axis that gives the best fit by comparing the adjusted R square of these multiple GLMs. A GLM with the largest adjusted R square and a positive regression coefficient for the grid signal regressor is selected. The reason we select the GLM with a positive regression coefficient is to avoid the inverted relationship between the hypothetical grid cell’s signal and the fMRI response (e.g. when movement is more aligned to the grid axis, the fMRI signal is lower). In rare case (<10% of voxels in our empirical data – see later sections) where all grid orientation models yield a negative regression coefficient, we simply select the GLM with the largest R square. To summarise, this iterative fitting process identifies which grid orientation best describes the fMRI signal in each voxel and in each scanning session.

Of note, we compared our numerical estimation method and the previous quadrature filter approach using simulated data in 2D (Supplementary Text 1). The simulation showed that both methods could detect a grid-like signal equally well at reasonable signal-to-noise ratios and sampling resolutions. We also tested whether the FCC and HCP grid models can be dissociated using synthetic data (Supplementary Text 2). This simulation showed that the correct model can be identified if the signal-to-noise ratio is high.

#### Testing for a grid signal in the fMRI data

We then test whether each voxel shows a consistent 3D grid signal across different scanning sessions by quantifying the regression coefficient of the grid signal model. For instance, if the fMRI data in one scanning session (e.g. run 1) is best fitted with a grid model that aligns at 15 degrees, we measure the grid score as the beta of the same grid orientation (15 degrees) model in the another scanning session (e.g. run 3). The beta values of voxels in the brain region of interest (ROI) are averaged for each participant, and a t-test is used to test whether the beta is positive at the group level (excluding outliers, participants with more than a standard deviation of 3 in our empirical data). This approach is similar to previous 2D grid analyses where the grid orientation is estimated from one half of the dataset and is tested on the other half of the dataset, and the regression coefficient is analysed at the group level (e.g. Doeller et al., 2010; a standard group level inference for fMRI experiments).

However, there is a difference between our study and some of the previous studies in terms of grid orientation averaging. In Doeller at al. (2010) and Horner et al. (2016), the estimated grid orientation of each voxel within the EC ROI was averaged, and this averaged grid orientation model was tested in the other half of the data. Here, we estimate and test the grid orientation model within each voxel, then we later summarise the grid score of voxels within the ROI. Neighbouring grid cells share a common grid orientation which is the essential property of grid cells that allows the detection of the direction-modulated signal at the fMRI voxel level, and earlier fMRI studies assumed one unique grid orientation for the entire EC. However, there is also evidence of multiple grid modules in the EC that have different grid orientations and scales (Stensola et al., 2012), and estimating and testing grid orientation at the voxel level, instead of the whole ROI, might maximise the sensitivity of analyses. This voxel-by-voxel estimation and test approach was used in a more recent 2D grid cell study (Nau et al., 2018).

### Experimental protocol

In this section we describe an empirical fMRI experiment where data were acquired while participants were exploring a virtual 3D environment. The experimental paradigm included a prescanning session with a head-mounted display. We applied our proposed 3D grid analysis to this dataset. This empirical dataset has been reported in a separate paper which investigated vertical and horizontal head direction encoding outside of the EC (https://www.biorxiv.org/content/early/2018/05/31/335976), and is summarised here for the reader’s convenience. The 3D grid code analysis reported in the current paper is novel and has not been published elsewhere.

#### Participants

Thirty healthy adults took part in the experiment (16 females; mean age 25.9±4.8 years; range 19-36 years; all right-handed). All had normal or corrected to normal vision and gave informed written consent to participation in accordance with the local research ethics committee.

#### The virtual environment

Participants were instructed that they were exploring a virtual zero-gravity environment, called “spaceship”, where they could move freely up, down, forwards and backwards. This spaceship had two rectangular compartments linked by a corridor (Fig. 6A, the environment can also be viewed in our online software). Participants could orient themselves in each compartment because the visual appearance of the walls differed (e.g. a window on the west side and a grey wall on the east side; the ceilings and floors also had different textures or hues). A snapshot of this virtual spaceship as seen from a participant’s perspective is shown in Fig. 6B,C. The virtual environment was implemented using Unity 5.4 (Unity Technologies, CA, United States).

**Fig. 6.**
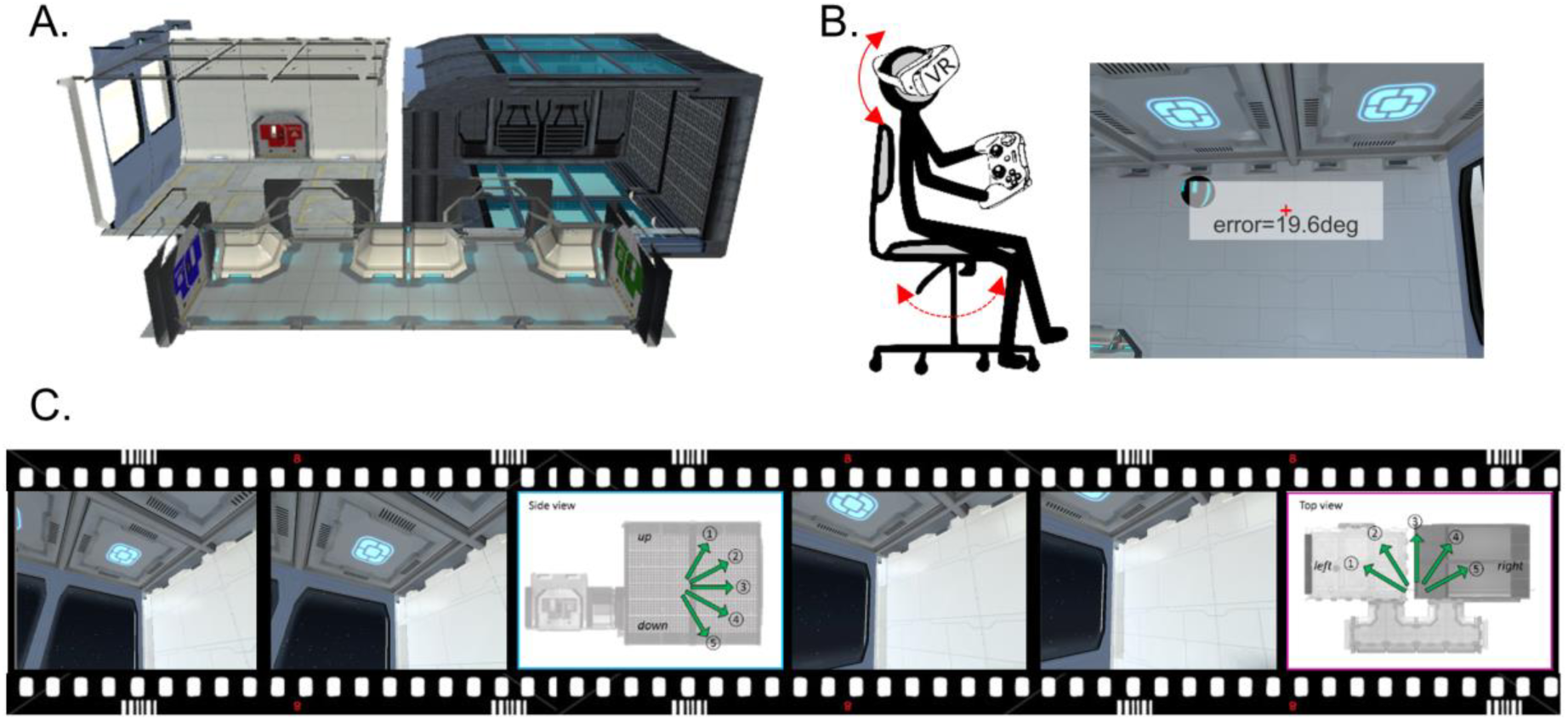
The empirical experiment. **A.** A side view showing the layout of the virtual 3D spaceship. **B.** Prior to scanning, participants explored the 3D environment while wearing a virtual reality head mounted display. **C.** During scanning, participants viewed a video rendered on a standard display. The video provided the feeling that participants were flying in a 3D trajectory within the spaceship. Participants occasionally indicated, via a keypad response, their vertical and horizontal movement direction when questioned.

The virtual spaceship was rendered on two different mediums for pre-scanning tasks and scanning tasks respectively: a head-mounted VR display (Samsung Gear VR, model: SM-R322 with Samsung Galaxy S6 phone) and a standard computer screen (Dell Optiplex 980 with an integrated graphic chipset). The VR display was used because it has been suggested that prior vestibular experience with a VR environment can later reinstate the relevant body-based cues during fMRI scanning, when only visual input is available due to head immobilisation (Shine et al., 2016). The VR head-mounted display provided participants with a fully immersive sensation of 3D space via its head motion tracking system, stereoscopic vision and a wide field-of-view (96°). A rotation movement in the VR display was made by a participant’s physical head rotation and a translational movement was made by a button press on the Bluetooth controller (SteelSeries Stratus XL, Denmark). For example, a participant could move up to the ceiling in the virtual spaceship by physically looking above and pressing the forward button. To rotate to the right, they physically rotated their head to the right or rotated their whole body when the required rotation was beyond the range of motion for neck rotation. For ease of rotation, participants were seated on a swivel chair.

During fMRI scanning, participants watched a video rendered on a standard computer screen (aspect ratio=4:3, Fig. 6C). The video was a first-person perspective that gave the participants the feeling of moving in a virtual spaceship. The stimuli were projected on the screen using a projector at the back of the MRI scanner bore (Epson EH-TW5900 projector), and participants saw the screen through a mirror attached to the head coil. The screen covered a field of view of ∼19° horizontally and ∼14° vertically.

#### Pre-scan: VR memory task

Wearing the VR display, participants freely explored the environment (5 minutes) and then performed a spatial memory test (15 minutes) where they had to recall the location of balls by physically directing their head to the remembered locations (Fig. 6). During the memory task, on each trial participants encoded the location of a floating ball by looking at it from different directions and distances for 18 seconds. Immediately after the encoding phase, participants were transported to a random fixed location and then had to look at the remembered location of the ball. They received feedback (angular error in degrees) after they made their decision. There were 16 main trials where participants had to locate the ball within the same compartment. We also added extra 6 trials where participants were asked to point a ball’s location across the wall from a different room. These trials were included to encourage participants to encode a global map of the environment.

#### fMRI scan: direction judgment task during passive viewing

During scanning, participants watched a video rendered on a standard display and performed a direction judgment task. The video provided participants with the feeling that they were flying in a controlled 3D trajectory within the spaceship (Fig. 6). The pre-programmed video allowed tight control of location, direction and timing for all participants. The trajectory consisted of multiple short linear movements (each of 3 seconds) followed by rotation (2/2.6 seconds). We restricted the range of movement directions (−60 to 60 degree with 30 degree steps, both vertically and horizontally, indicated by arrows in Fig. 6C) to increase the frequency of each movement direction visited within the limited scanning duration. All participants followed the same trajectory without abrupt rotations where each of 25 directions (5 levels of pitch x 5 levels of azimuth) was evenly sampled (min = 18, max = 20 trials). A constant linear and angular velocity was applied in order to control velocity, which can also influence grid cells’ activity (Sargolini et al., 2006). If a participant reached the boundary of the spaceship, a blank screen appeared for 2 seconds and then a trajectory started again from the other end of the spaceship. For 25% of the time, a question screen appeared immediately after a linear movement and participants indicated the direction of their last movement by pressing an MR-compatible button pad (a 5-alternative forced choice question with a time limit of judgment task was used to ensure participants kept track of their movements during scanning. The two compartments of the spaceship were visited alternatively for each of 4 scanning sessions. Half of the participants started in one compartment and half started in the other compartment. Each scanning session lasted ∼11 minutes with a short break between the sessions, making a total functional scanning time of ∼50 min.

#### Post-scan debriefing

After scanning, participants were debriefed about how much they felt immersed in the virtual environment during the pre-scan session with VR head mounted display and during scanning. Participants chose from multiple options: “I felt like I was really in the spaceship”; “I occasionally thought about the environment as being computer-generated, but overall the environment was convincing and I felt I was moving around in the spaceship”; “I was often distracted by the feeling that I was not in a real environment”.

#### Behavioural analyses

For the pre-scan memory task, we report the mean angular deviation for the main trials (where participants had to look towards the remembered location of the ball within the same compartment of the environment). We then report the overall accuracy of the direction judgment task during scanning (chance=20%) to check whether participants knew in which direction they were moving in the 3D environment.Further data and analyses related to the vertical and horizontal direction sensitivity, are available in preprint form here (https://www.biorxiv.org/content/early/2018/05/31/335976). For the debriefing question, we counted the number of responses for each option.

#### Scanning and pre-processing

T2*-weighted echo planar images (EPI) were acquired using a 3T Siemens Trio scanner (Siemens, Erlangen, Germany) with a 32-channel head coil. Scanning parameters optimised for reducing susceptibility-induced signal loss in areas near the orbitofrontal cortex and medial temporal lobe were used: 44 transverse slices angled at −30°, TR=3.08 s, TE=30 ms, resolution=3×3×3mm, matrix size=64×74, z-shim gradient moment of −0.4mT/m ms (Weiskopf et al. 2006). Fieldmaps were acquired with a standard manufacturer’s double echo gradient echo field map sequence (short TE=10 ms, long TE=12.46 ms, 64 axial slices with 2 mm thickness and 1 mm gap yielding whole brain coverage; in-plane resolution 3 x 3 mm). After the functional scans, a 3D MDEFT structural scan was obtained with 1mm isotropic resolution.

Preprocessing of data was performed using SPM12 (www.fil.ion.ucl.ac.uk/spm). The first 5 volumes from each functional session were discarded to allow for T1 equilibration effects. The remaining functional images were realigned to the first volume of each run and geometric distortion was corrected by the SPM unwarp function using the fieldmaps. Each participant’s anatomical image was then coregistered to the distortion corrected mean functional images. Functional images were normalised to MNI space and smoothed with 6mm kernel.

#### ROI selection

We used anatomical ROIs, left and right EC masks (Fig. 7). The ROIs were manually delineated on the group-averaged MRI scans from a previous independent study on 3D space representation (Kim et al., 2017) following the protocol in Pruessner et al. (2002). The number of functional voxels (3 x 3 x 3 mm) within the ROI masks was 47 (left) and 49 (right). Of note, EC was further divided into posterior medial and anterior lateral parts in one previous fMRI study (Bellmund et al., 2016), based on the finding in rodents that grid cells are typically reported in the medial EC. However, our study used standard resolution fMRI and further segmentation of this kind was not feasible. Functional specialisation within the EC is an interesting topic that needs to be further addressed in future studies with high-resolution scanning sequences.

**Fig. 7.**
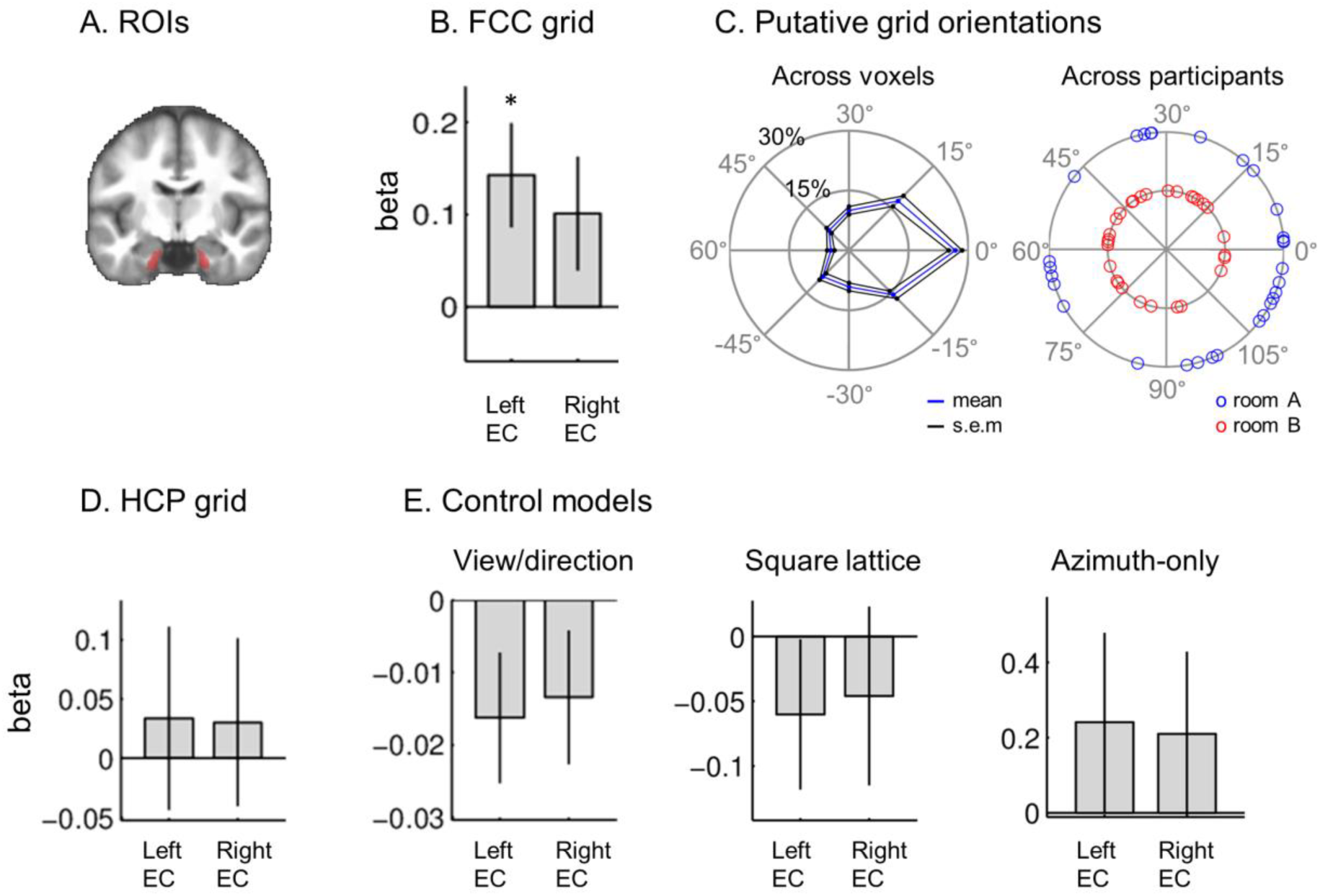
3D grid-like representations in the left EC. **A.** Bilateral EC (red) ROIs are shown on the group-averaged structural MRI scan. **B.** The mean beta of the FCC grid cell model was significantly positive in the left EC. There was a similar trend in the right EC (p=0.054). **C.** Putative FCC grid orientation in the left EC was clustered around the mean grid orientation within participants (the left rose plot), but random across participants (the right polar plot). **D.** The HCP model was not significant in either EC (see the main text for a methodological limitation related to this). **E.** Control models (view or direction encoding models, a square lattice model and an azimuth-only grid model) were also not significant in the EC. *p<0.01. The error bars are 1 SEM.

Another important consideration for selecting EC ROIs is that EC is notoriously difficult to image because of fMRI susceptibility artefact in this vicinity. Although sequence development continues in this regard, EC still has inherently low raw BOLD signal compared to other cortical regions. Crucially, standard fMRI analysis software like SPM excludes voxels of low signal by default. The “global masking threshold” parameter in the first-level model specification in SPM determines which voxels are to be included in the analysis based on the raw intensity, and EC voxels can often be excluded. It can also result in a different number of voxels in an EC ROI for each participant. We caution researchers about excluding voxels for two reasons. First, an exclusion criterion based on the mean BOLD intensity can be arbitrary. Depending on the version of the software, some voxels can be included or excluded from the analysis. Second, raw BOLD intensity alone does not predict whether a voxel shows functional modulation. For instance, whereas the raw signal intensity of cerebrospinal fluid (CSF) is higher than most other cortical areas, we rarely observe meaningful signals in the CSF in typical cognitive experimental paradigms. In our study, we defined the EC ROIs purely anatomically, without excluding any voxels based on raw signal intensity.

#### Main grid analysis

We applied the 3D grid analysis that we described in detail above to the preprocessed fMRI dataset. In essence, we estimated the orientation of the 3D grid axis for the FCC model in each voxel and scanning session by iteratively fitting the fMRI time series to the predicted grid alignment score defined as the cosine of movement direction and the nearest grid axis. The grid model was tested on another scanning session. Because our virtual spaceship had two compartments, we trained and tested the grid cell models within each compartment, and averaged the regression coefficient of the two compartments.

We then tested whether the estimated grid orientation of the FCC model was clustered across voxels within participants or across multiple participants. To test the voxel-wise clustering within participants, we calculated the percentage distribution of angular distance between the circular mean grid orientation and the estimated grid orientation of each voxel within ROIs in each participant (=histogram with bins centred at −45, −30, −15, 0, 15, 30, 45, 60 degree difference in putative grid orientation). If the grid orientation was clustered across voxels, this distribution would be non-uniformly distributed with the mode centred at 0. We averaged the angular distance distribution across participants, then applied a V-test for non-uniformity. To test the clustering across participants, we applied a Rayleigh test for non-uniformity to the circular mean grid orientation across participants. If the grid axis was anchored to visual features in the environment such as landmarks or the boundary, every participant would exhibit a similar grid orientation. The circular mean and non-uniformity test was computed using the CircStat2012a toolbox (Berens, 2009). We normalised the grid orientation into 2*pi radians before we applied the V-test or Rayleigh test because the putative grid orientation was defined between 0 to 120 degrees whereas standard circular statistics is applied for 0 to 360 degrees.

We also tested whether the HCP grid model explained signal in the EC despite the methodological limitation which we explained earlier.

#### Control analysis – direction or view encoding model

Our 3D grid analysis (as well as the 2D grid analyses in the literature) relies on the dependency of the neural signal on movement direction, and one concern is whether a neural signal that is responsive to one particular direction (or the view associated with a direction) could be weakly correlated with a grid model and so identified as a grid signal. This was why we used a direction (or view) encoding model as a control analysis. We created a direction-sensitive model signal which was sensitive to one of nine 3D directions that were visited by participants in a virtual environment. The nine directions were regularly sampled both horizontally and vertically: (azimuth, pitch in degrees) = (−60, −60), (−60, 0), (−60, 60), (0, −60), (0, 0), (0, 60), (60, −60), (60, 0), (60, 60). Following Bellmund et al. (2016), we assumed that each direction-sensitive neural response had a margin of 30 degrees. This meant that neurons or voxels that responded strongly to (0, 0) direction would also respond strongly to (±30, ±30), and would respond weakly to the rest of the movement directions. We convolved the binary direction response vector with the hemodynamic response function. We created a GLM similar to the grid cell model described in the previous section but now the grid signal was replaced by the direction encoding signal. Again, the best direction-encoding model was selected for each voxel from one scanning session and then tested on a different scanning session. If voxels in our ROIs (left and right EC) responded to unique directions, we would see a significantly positive regression coefficient for a direction model at the group level.

#### Control analyses – other grid models

In 2D, a non-hexagonal grid model, such as a 4-fold symmetry, has been used as a control model (Doeller et al. 2010). Similarly, we tested whether fMRI signal in the EC was explained by a square lattice model (Fig. 8). A square lattice model has a lower packing density than the FCC and HCP models, so the square lattice model is not an optimal way of encoding 3D space. Just as in our testing of the FCC and HCP models, we assumed that the activity of a grid voxel was modulated by the alignment score (cosine of angle) between movement direction and the grid axis orientation (Fig. 8).

**Fig. 8.**
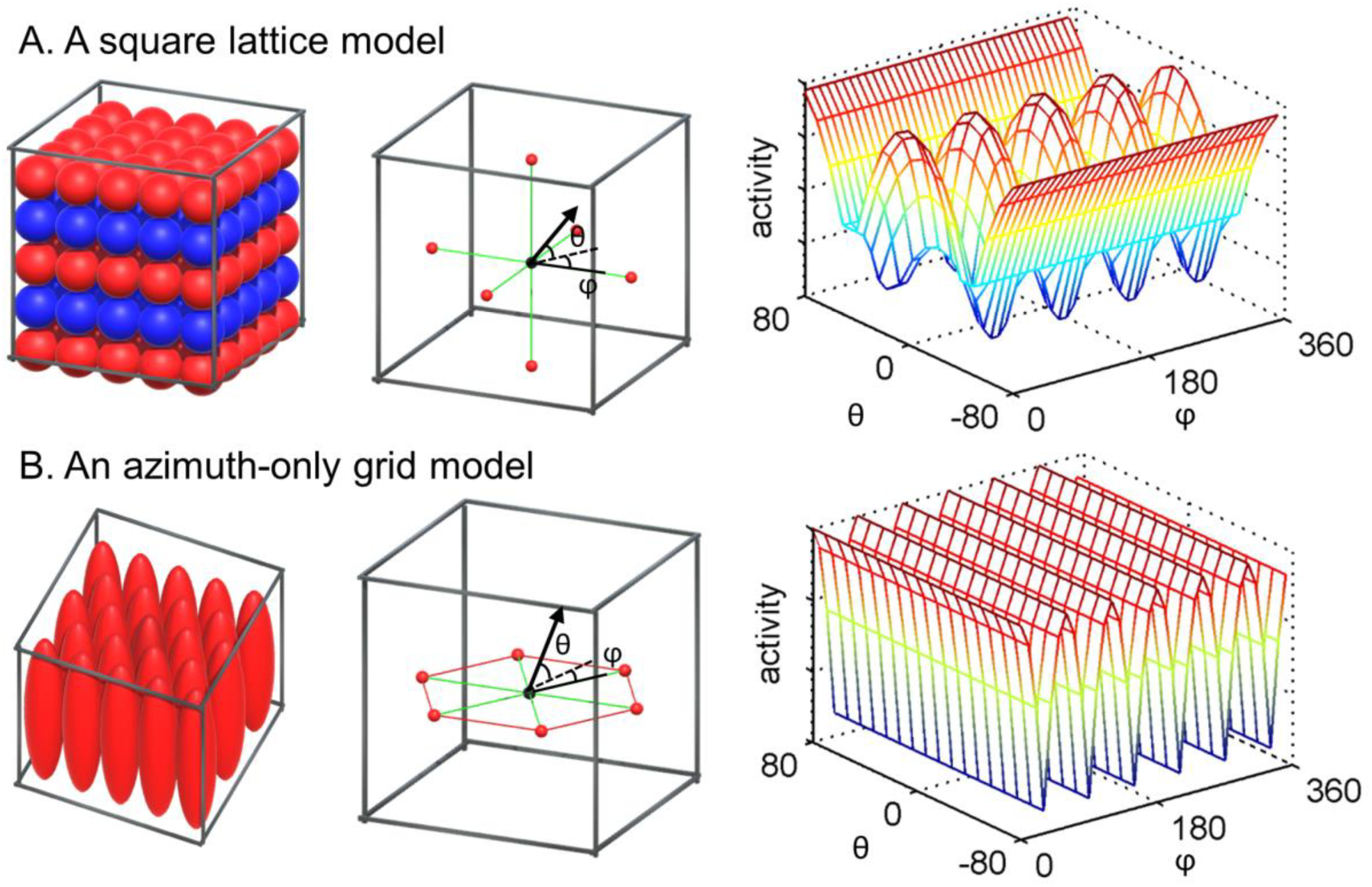
Control grid models. **A.** Receptive fields of hypothetical grid cells which follow a square lattice arrangement (the left panel). The grid activity is expected to be modulated by the movement direction (the black arrow) relative to the orthogonal grid axis (the green lines, the middle panel). It displays 90° periodicity for azimuth (ϕ) when vertical pitch (θ) is close to zero (the right panel). **B.** Receptive fields of hypothetical grid cells which show hexagonal periodicity only along the horizontal axes and not along the vertical axis (the left panel). The grid activity is expected to be modulated by the horizontal movement (ϕ) independent of whether a participant is moving up or down (θ, the middle and the right panels).

We also tested a hexagonal grid model which was only responsive to the horizontal axis (“Azimuth-only”). This model is related to the previous observation in rats that receptive fields of grid cells were vertically elongated in a spiral staircase apparatus (Hayman et al. 2011) (Fig. 8). We know nothing about whether or not there are direction-modulated signals when grid cells have such anisotropic receptive fields, but we think it is reasonable to assume that grid activity would be only modulated by the horizontal angle between movement direction and the grid axis, independent of pitch, in this case (Fig. 8). For example, grid activity would be high if a participant moves in a 0°, 60°, 120°, 180°, 240°, 300° direction (azimuth-wise) independent of whether they move up or down.

#### Control analysis –primary sensory cortex ROIs

We tested the FCC grid model in size-matched control brain regions to reassure that the grid-like representation we identified in EC was not merely an artefact that was present across the whole brain. We defined spherical ROIs (radius 7mm) centred at peak coordinates from the fMRI meta-analysis toolbox Neurosynth using the key words “primary auditory” [−44, −24, 8] and “primary visual” [−8, −86, 0] (Yarkoni et al. 2011). In addition, we tested the view encoding model in the primary visual cortex ROI.

## Empirical results

### Behavioural

During the pre-scan memory task, participants were reasonably successful at locating a ball’s position (mean error 21°, SD 9°, Fig. 6B). The mean accuracy of direction judgments during scanning was well above chance level (mean 74%, SD 16%; chance 20%), suggesting that participants knew their 3D movement direction in the virtual environment during scanning.

The rating data showed that participants felt immersed in the virtual environment, with the vast majority choosing either “I felt like I was really in the spaceship” (57% for the pre-scan VR head mounted display task, 10% for the scanning task) or “I occasionally thought about the environment as being on a computer screen, but overall the environment was convincing and I felt I was moving around in the spaceship” (43% for the pre-scan VR head mounted display task, 80% for the scanning task). This result implies that our virtual environment effectively conveyed a sense of being in 3D space.

### fMRI – main grid analysis

We tested whether fMRI signals in the left and right EC was modulated by participants’ 3D direction as predicted by the FCC lattice model. The FCC grid model was significant in the left EC (t(28)=2.6, p=0.008, one-sided), and showed a trend in the right EC (t(28)=1.7, p=0.054, one-sided) (Fig. 7).

The putative FCC grid orientation of each voxel was clustered around the mean orientation within participants (V-test in the left EC, V=33.6, p=1.0*10^-6^; the right EC, V=30.5, p=7.9*10^-6^, Fig. 7C) similar to previous studies (Nau et al., 2018; Julian et al., 2018). By contrast, the mean grid orientation was not significantly clustered across participants (Rayleigh’s test for non-uniformity in the left EC, room A, z=1.6, p=0.2; room B, z=1.5, p=0.2; the right EC, room A, z=1.6, p=0.2; room B, z=0.3, p=0.8, Fig. 7C), suggesting that the grid axis was not anchored to, or driven by, particular features of the environment. This non-clustered grid orientation across participants is consistent with previous studies which used a circular arena (Doeller et al., 2010; Nau et al., 2018). Of note, Julian et al. (2018) observed a clustering of grid orientation around 7.5 degrees to the cardinal axis (modulo 15 degree) for a square arena, but not for a rectangular environment that was similar to that of our spaceship.

We also tested the HCP grid model with the assumption that the fMRI signal is modulated by the local grid axis. The HCP model did not significantly explain the response of either EC (the left EC, t(29)=0.4, p=0.3, the right EC, t(29)=0.4, p=0.3, one-sided, Fig. 7D). However, as we explained in the Methods section, an fMRI analysis that relies on a direction-modulated grid signal is not ideal for detecting the HCP model because it lacks a global grid axis. Thus, a further comparison between the FCC and HCP models requires future electrophysiological studies that can directly assess the receptive fields of grid cells.

### fMRI – control analyses

To exclude the possibility of a neural signal sensitive to one particular direction (or associated view) being identified as a grid voxel, we tested a unique direction encoding model as a control. The direction encoding model was not significant in either EC (left EC, t(29)=-1.8, p=0.9; right EC, t(29)=-1.5, p=0.9, one-sided, Fig. 7E), suggesting that the FCC grid-like signal that we observed in the EC was not driven by one particular direction.

We also tested a square lattice model and an azimuth-only model where vertical pitch was ignored. Neither of these models significantly explained the fMRI signal in the EC (square lattice model: left EC, t(29)=-1.1, p=0.9; right EC, t(29)=-0.7, p=0.7; azimuth-only model: left EC, t(29)=1.0, p=0.2; right EC, t(29)=1.0, p=0.2, one-sided, Fig. 7E).

Finally, we tested the FCC grid model in size-matched primary auditory cortex and visual cortex ROIs. The primary auditory cortex did not show a grid-like signal (primary auditory cortex, t(29)=0.6, p=0.6, one-sided), suggesting that the FCC grid-like signal that we identified in the EC was not a spurious effect that was detectable anywhere in the brain. However, the FCC grid model was significant in the primary visual cortex ROI (t(29)=1.71, p=0.049, one-sided) (Supplementary Fig. 3). We think the grid model partly explained this response because the primary visual cortex is modulated by views and the view-dependent signal could be weakly correlated with the direction-modulated grid model. Indeed, our view encoding control model was highly significant in the primary visual cortex (t(29)=7.35,=2.1*10^-8^, one-sided) and R square was higher for the view encoding model than the FCC grid model in the visual cortex, implying that this response was better explained by the view encoding model than the FCC grid model (Supplementary Fig. 3). This result was in contrast to the response in EC where only the FCC grid model was significant, not the individual view encoding model. If there were periodic visual features and these were the sole reason for the 3D grid-like signal observed in the EC then the response profiles should have been identical for the visual cortex and the EC, which they were not.

## Discussion

In the present study, we presented a novel analysis method to investigate 3D grid codes noninvasively in humans. Simulation and actual fMRI data suggested that it is possible to probe one type of 3D grid model, an FCC lattice model, by relying on direction-modulated grid signals at the macroscopic level. We also developed associated software to help researchers visualize 3D receptive fields of grid cells and predict their responses. Here we discuss the implications and limitations of our study and make suggestions for future studies on 3D grid codes.

The main finding of this study relates to our probing of putative grid cells using fMRI by predicting the neural signal as a function of 3D movement direction and the grid axis. The principle of measuring direction-modulated grid signals has been widely used in 2D (Doeller et al., 2010; Constantinescu et al., 2016; Bellmund et al., 2016). However, in our study we extended, for the first time, this principle into 3D volumetric space, thereby opening up the possibility of empirically studying grid cells in high-dimensional space. We successfully demonstrated the feasibility of this analysis approach by finding our data was concordant with the FCC model in the left EC, the candidate brain structure for 3D grid encoding. Importantly, we also exposed the fundamental limitation of movement direction-based grid analysis in 3D. Unlike in typical 2D environments where grid fields align with regular grid axes (but see Krupic et al., 2014 for distorted grid axes in 2D), some proposed 3D grid models like the HCP model lack global grid axes (Mathis et al., 2015). The summed response of numerous grid cells is unknown in the absence of global axes. Nevertheless, we attempted to test the HCP model using locally defined grid axes, and the HCP model did not fit our empirical data. Due to the grid axes issue, we cannot claim superiority of the FCC model over the HCP model. Rather, we suggest that direct recording of grid cells is needed to compare different potential 3D grid models, as they can circumvent the issue of grid axis and direction-modulation.

Once there is a fuller understanding of the cellular physiology of grid cells, it will be possible to determine the optimal fMRI analysis protocol by considering the multiple factors we have described here, such as whether the orientation of the 3D grid axis is parallel to the ground or not, the precise model between the grid alignment and the fMRI signal (e.g. cosine, linear, binary), and the distribution of the grid orientation across different voxels within EC. Furthermore, a better understanding of the spatial organisation of grid cells might allow us to measure the grid signal without relying on the direction-modulation principle. For example, though speculative, there might be a bias in grid phase at the voxel level whereby some voxels contain more grid cells with a particular grid phase. This would result in a periodic response as a function of location which may be detectable by a spectral analysis. This might enable us to directly compare the FCC, HCP and other 3D grid models. Previous optical imaging methods revealed a micro-organisation of 2D grid cells in the medial EC, although the spatial scale was much finer than the typical fMRI voxel size (Heys et al., 2014; Gu et al., 2018).

Regarding the experimental design for future investigations of 3D grid code, the use of an immersive volumetric environment is important. A 3D lattice structure like FCC is optimal for encoding volumetric space (Mathis et al., 2015), and the existence of substructures like 2D walls or a 1D track could affect the response of grid cells. For instance, grid cells recorded in rats moving on a sloped terrain showed a firing pattern similar to a 2D horizontal plane rather than a 3D lattice pattern (Hayman et al., 2015). Therefore, in the current study we built a fully volumetric virtual “zero-gravity” environment where participants could move freely in all directions. Furthermore, we complemented the fMRI scanning, where only visual input was available because of in-scanner head immobilisation, by using a VR head mounted display during pre-scan tasks as in Shine et al. (2016). We believe this pre-scan experience of physical head rotation when using the VR head mounted display was particularly helpful to participants in building the mental and neural representation of a 3D space, and this was supported by participants’ reports in the debriefing session at the end of the experiment.

Future studies examining 3D grid codes should also consider using active movement paradigms that sample all possible movement directions. In the current study, participants were passively moved during scanning for ease of controlling movement trajectories and to achieve even sampling of each 3D direction. Although previous fMRI studies have successfully observed grid signals using imagination tasks without active self-motion (Bellmund et al., 2016; Horner et al., 2016), it is known that passive movement disrupts velocity-modulated theta and grid firing in rodents (Winter et al., 2015). Self-motion signals are therefore critical for path integration associated with grid cells (McNaughton et al. 2006) and, consequently, more robust grid signals might be detected if active (virtual) movement paradigms are employed. One might also improve the power of detecting grid signals by sampling all 3D directions. In the present experiment, we only sampled movement directions spanning 120° vertically and horizontally due to limited scanning time. This limited sampling caused unbalanced data when investigating individual directional responses relative to the grid axis (e.g. participant A moved 0° to 120° relative to the putative grid orientation whereas participant B moved −30° to 90° relative to the grid orientation).

In summary, we believe that our experimental paradigm, analysis method and software serve as a useful initial stepping-stone for studying grid cells in realistic 3D worlds. Animal electrophysiology and human fMRI studies also suggest that a grid code is employed to encode not only physical space, but also more abstract knowledge (Aronov et al., 2017; Constantinescu et al., 2016; Nau et al., 2018; Julian et al., 2018), and we hope our approach will in due course also encourage interrogation of abstract high-dimensional cognitive processes.

## Acknowledgements

This work was supported by the Wellcome Trust (101759/Z/13/Z to E.A.M.; 203147/Z/16/Z to the Centre; 102263/Z/13/Z to M.K.) and a Samsung Scholarship (to M.K). We thank Kate Jeffery, Tim Behrens, Alexa Constantinescu and Neil Burgess for helpful discussions, Anna Monk for drawing Figure 6B, and Marshall Dalton for his advice on neuroanatomy.

The authors declare no competing financial interests.

---

**Supplementary Fig. 1.**
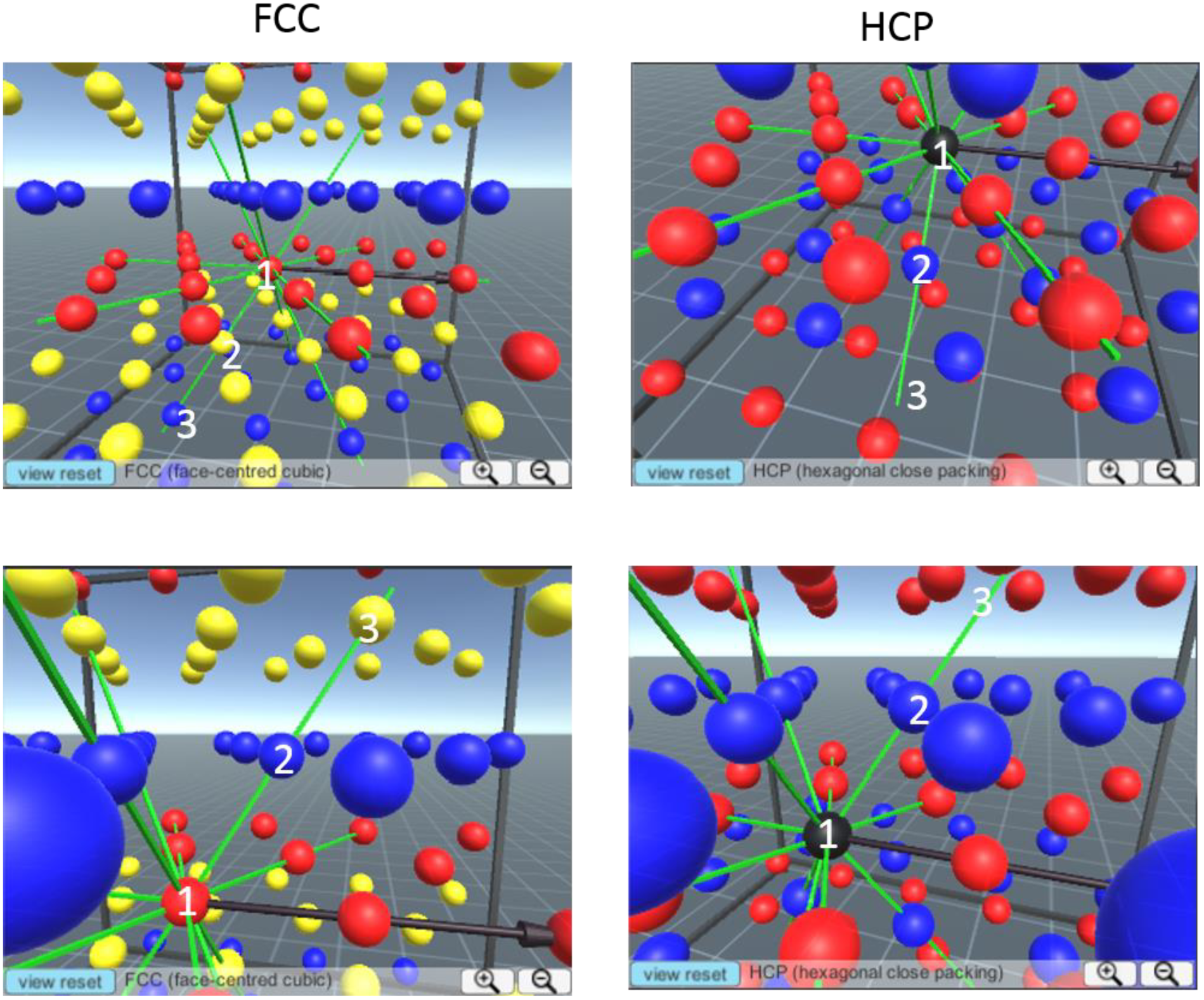
Grid axes of the FCC and HCP arrangements. FCC is a pure lattice with centre-symmetry. If one moves along the main grid axis from one node, every grid field will be passed (three nodes labelled as ‘1’, ‘2’, ‘3’ in this example). In contrast, HCP does not have such symmetry. When one moves in the locally-defined grid axis, the grid field at the next layer is not passed (‘3’ does not pass the centre of the grid fields). It is easier to appreciate this 3D arrangement by using our visualization software (www.fil.ion.ucl.ac.uk/Maguire/grid3D_gui).

**Supplementary Fig. 2.**
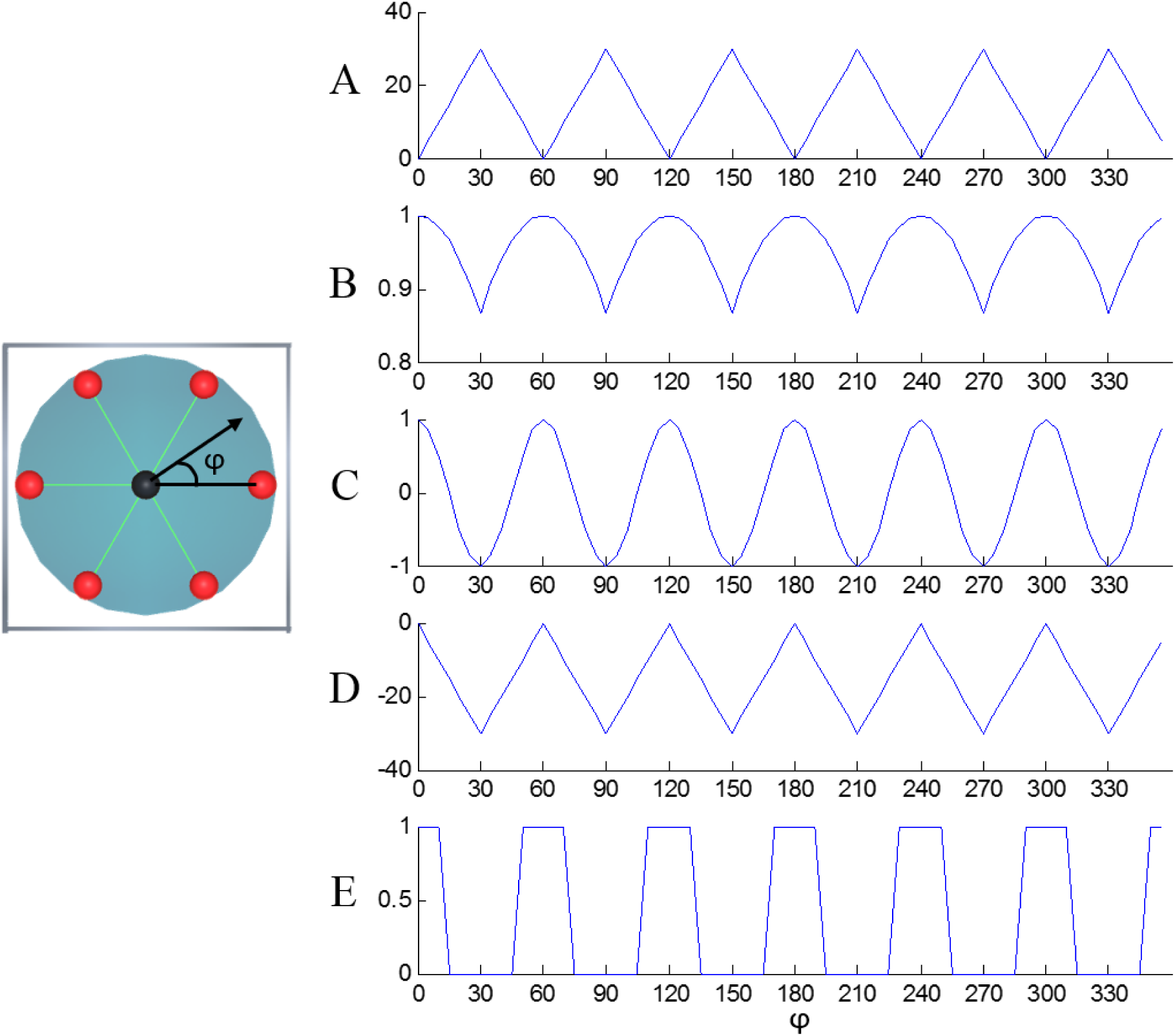
Possible relationships between grid alignment and the fMRI signal. **A.** An angle between the movement direction (black arrow) and the nearest grid axis (green lines). **B.** A cosine of the nearest angle. **C.** A sinusoidal signal with a periodicity of 60 degrees. **D.** A linear signal (the negative of the nearest angle). **E.** A binary signal.

**Supplementary Fig. 3.**
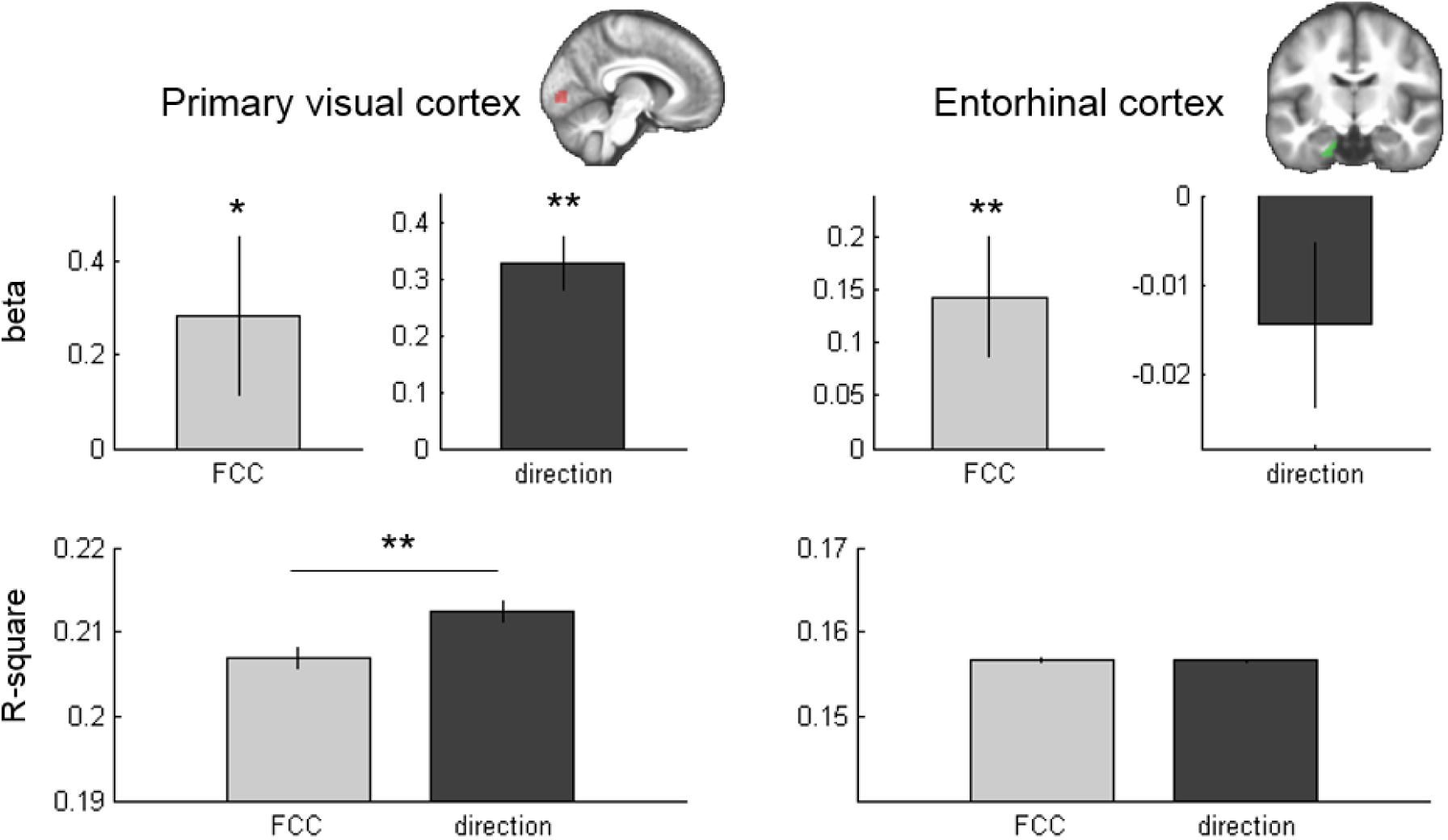
Comparison of primary visual cortex and the left entorhinal cortex. For ease of comparison, on the left we reprise the entorhinal cortex result from the main text. The FCC grid model and the direction model were tested in primary visual cortex and the left entorhinal cortex. In the visual cortex, both the FCC and direction encoding models were significant. However, the FCC model fitted better than the direction encoding model (larger R-square). In contrast, only the FCC model was significant in the entorhinal cortex. * p<0.05, **p<0.01.

## Supplementary Text 1

Comparison of numerical estimation and quadrature filter approaches in simulated 2D grid signals.

### Introduction

In this simulation, we generated synthetic 2D grid signals to test whether our numerical grid orientation estimation method can detect such signals. We compared how well the numerical estimation methods with different sampling resolutions and a previous analytic approach (the quadrature filter, e.g. Doeller et al. 2010) detected the grid signal in the presence of noise.

### Methods

We first created two virtual direction trajectories (approximating two scanning sessions of 10 minutes length each) where the range of movement (θ) was restricted to [-60°, 60°], similar to our empirical data (see Supplementary Fig. 4 below). Next, we computed the grid alignment score vector by assigning random grid orientations (ϕ) for 30 virtual participants using the formula, cos (6* (θ – ϕ)). This hypothetical grid cell activity was convolved with a canonical hemodynamic response function and then Gaussian noise of varying size (SNR=1, 0.1, 0.01) was added.

In the numerical estimation method, we iteratively fitted the hypothetical grid signals when different grid orientations were assumed (either 15°, 7.5° or 1° resolution) and selected the grid orientation which had the best fit to the actual data in each scanning session (see details in the main text). We tested whether the grid model selected in the first session resulted in a positive regression coefficient in the second session and vice versa (a one-sided t-test). For the quadrature filter approach, we estimated the grid orientation (ϕ) using the regression coefficient of cos(6*θ) and sin (6*θ) regressors, ϕ=arctan(b1/b2) for each scanning session. We tested whether a new cosine regressor with the grid orientation estimated from the first session resulted in a positive regression coefficient in the second session and vice versa.

### Results

Supplementary Fig. 5 below shows group mean grid scores (beta) and R-square values for varying SNR. All numerical estimation methods and quadrature filter approaches could reliably detect the grid signal when SNR was high (Supplementary Fig. 5A, 15°, t(29)=33.5, p<0.001; 7.5°, t(29)=55.0, p<0.001; 1°, t(29)=97.3, p<0.001, quadrature, t(29)=98.1, p<0.001) or middle (Supplementary Fig. 5B, 15°, t(29)=5.3, p<0.001; 7.5°, t(29)=4.9, p<0.001; 1°, t(29)=5.0, p<0.001, quadrature, t(29)=5.1, p<0.001). None of the methods could detect the grid signal if the SNR was low (Supplementary Fig. 5C, 15°, t(29)=0.0, p=0.5; 7.5°, t(29)=-0.2, p=0.6; 1°, t(29)=-0.2, p=0.6, quadrature, t(29)=-0.2, p=0.6).

When SNR was high, the quadrature filter approach and numerical method with fine sampling resolution (1°, which virtually converges to an analytic solution) was better at detecting a grid signal compared to the numerical method with coarser sampling resolution (7.5°, 15°), in terms of a higher grid score and R-square (Supplementary Fig. 5A, grid score, F(3,87)=18.8, p<0.001; R-square, F(3,87)=27.9, p<0.001). This was due to discretized estimates of the grid orientation when coarser sampling resolution was used.

The numerical methods with varying sampling resolution and the quadrature filter approach showed similar grid scores and R-square values when SNR was middle (Supplementary Fig. 5B, grid score, F(3,87)=1.3, p=0.3; R-square, F(3,87)=1.0, p=0.4) or low (Supplementary Fig. 5C, grid score, F(3,87)=0.7, p=0.6; R-square, F(3,87)=2.2, p=0.1).

### Discussion

Both analytic methods and numerical methods could detect underlying grid signals if the noise level was not too high. The analytic method and a numerical method with fine sampling resolution performed better than a numerical method with coarse sampling resolution if SNR was high. Real fMRI data is often noisy (e.g. the adjusted R-square value of our actual data in the entorhinal cortex was only 0.15) and the sampling resolution of the numerical method might have a negligible effect on detecting grid signal. Indeed, FCC grid scores in our actual fMRI data were comparable when we estimated the grid orientation with either 15° or 7.5° resolution.

---

**Supplementary Fig. 4.**
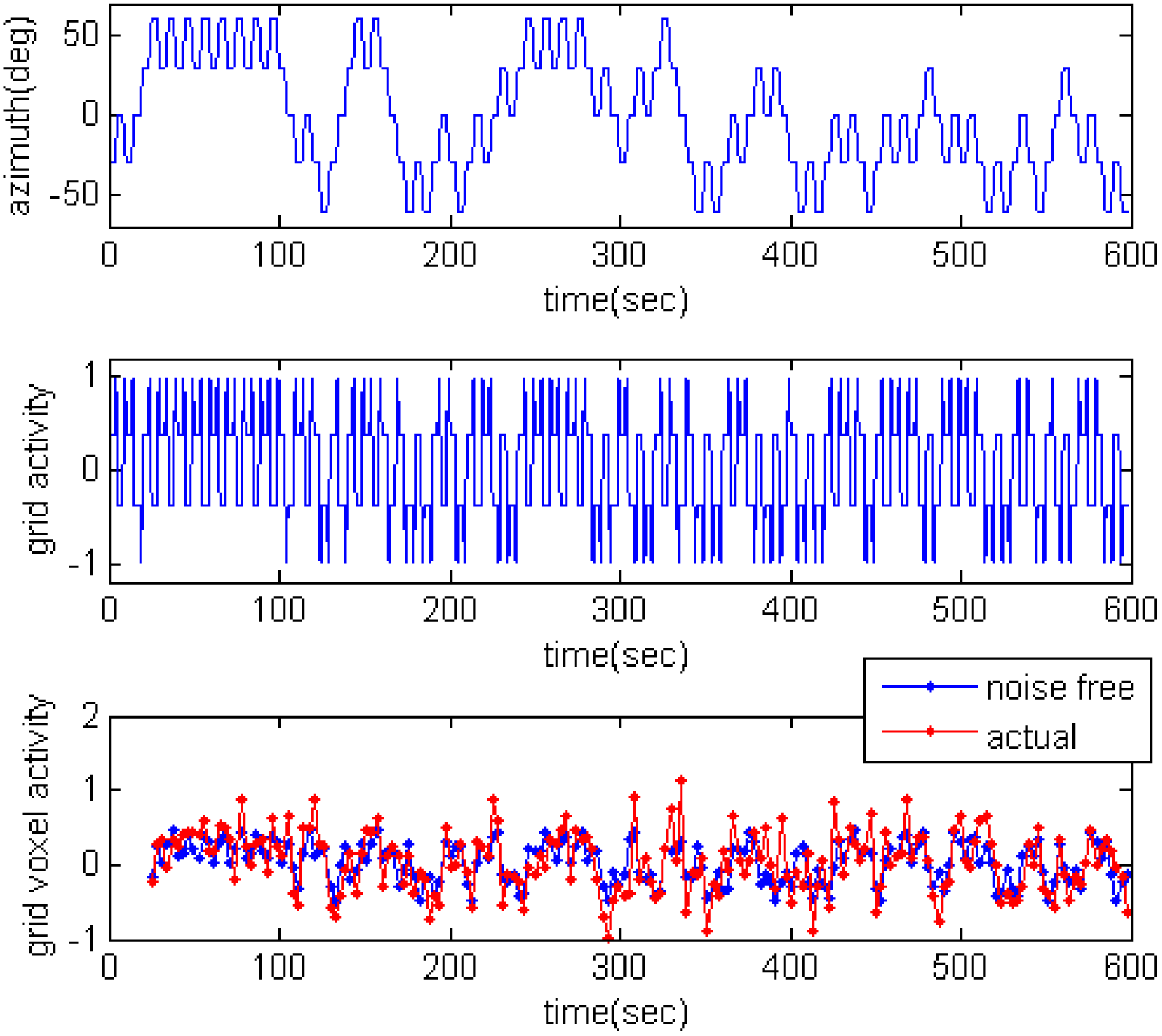
Simulated grid cell activity. Top panel, virtual movement trajectory of a participant. Middle panel, hypothetical grid cell activity of one participant whose grid orientation (ϕ) was 41°. Bottom panel, hypothetical grid cell activity was convolved with a hemodynamic response function, resulting in grid voxel activity. Before (“noise free”) and after (“actual”) the addition of Gaussian noise (SNR=1).

**Supplementary Fig. 5.**
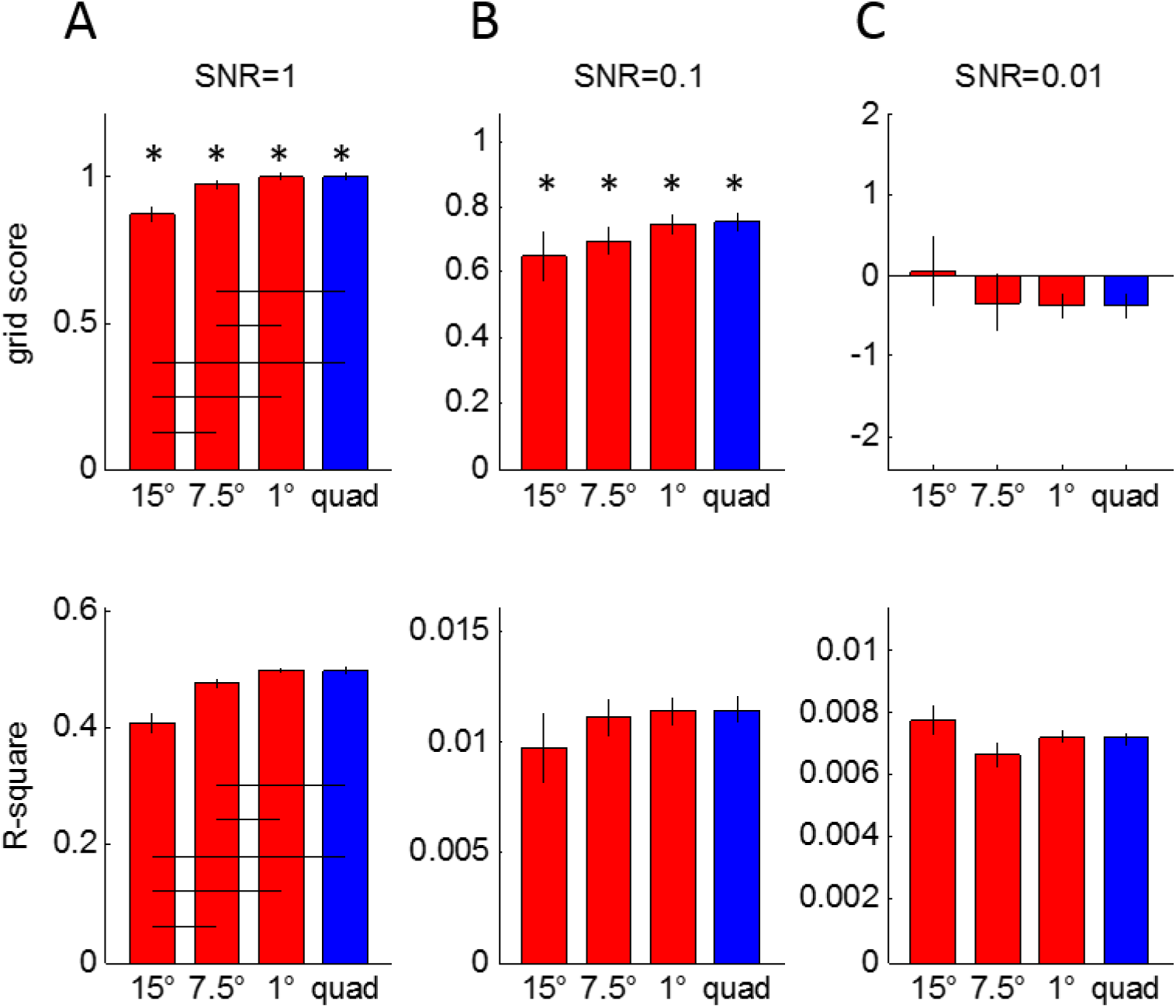
Comparison of the numerical estimation and quadrature filter approaches in 2D simulated data. When SNR was relatively high (**A** and **B**), both numerical and filter approaches correctly revealed the grid signal (positive grid score). When SNR was low (**C**), neither of the methods recovered the grid signal. Quadrature filter approaches or the numerical estimation with fine resolution (1°) were more sensitive to detecting grid-like signals than numerical estimation methods with coarse sampling resolutions (15°, 7.5°), only when SNR was high (**A**). All methods yielded similar results when SNR was middle or low (**B** and **C**). *p<0.05. Black horizontal bars indicate a significant difference between the conditions (post-hoc pairwise t-tests). Error bars are standard error adjusted for a within-subjects design.

## Supplementary Text 2

Comparison of the FCC and HCP models in simulated 3D grid signals.

### Introduction

In this simulation, we generated synthetic 3D grid signals to test whether our numerical grid orientation estimation method can correctly identify the FCC and HCP models. We already discussed the challenge of detecting the HCP model due to its absence of global grid axes (Methods section in the main manuscript), and for the purpose of comparing the two models, we assumed here that the HCP model is modulated by locally defined grid axes. The goal of this simulation was to investigate whether the two models could be distinguished despite their similarities and the presence of noise when limited movement directions were sampled.

### Methods

We used virtual 3D trajectories equivalent to those used in the actual experiment (from −60 to 60 degrees vertically and horizontally, ∼240 TRs for each scanning session) (Supplementary Fig. 6A). We assigned random grid orientations for 30 virtual participants (0, 7.5, 15, 22.5, …, 120 degrees) and created a direction-modulated grid signal that followed either the FCC or HCP model (=cosine of the angle between the movement directions and the nearest grid axis) (Supplementary Fig. 6B). We reiterate that the exact form of the direction-modulated signal is unknown in the HCP arrangement because of the lack of a global grid axis, and here we assumed that the HCP signal would be dependent on the local grid axes. A canonical hemodynamic response function was applied and Gaussian noise was added to the hypothetical grid signal (SNR=1 or 0.05).

We estimated the grid orientation of each session by iteratively fitting the hypothetical grid signals, separately for the FCC or HCP models with 15° resolution. We tested whether a regression coefficient (grid score) for the selected grid model was positive at the group level (a one-sided t-test). We compared the grid score and R square between the FCC and HCP models when the true data were generated with either the FCC or HCP models.

### Results

When the SNR was high (SNR=1), the FCC model showed a higher grid score and larger R squared than the HCP model when the true model was the FCC (Supplementary Fig. 6C), and vice versa when the true model was the HCP (Supplementary Fig. 6D). This means that the two models were correctly and unequivocally distinguished. Of note, grid scores were significantly positive for both FCC and HCP models regardless of whether the true model was FCC or HCP. This was due to the similarity between the FCC and HCP alignments (e.g. 9 out of 12 neighbouring fields were at the same 3D locations for the two alignments, Figure 2 in the manuscript).

When the SNR was low (0.05), we repeated the simulation 100 times by changing the random number generator setting (“rng” function in MATLAB) to avoid the risk of reporting an extreme case in this low SNR regime. Here, we describe the cases when the true model was FCC (the results were similar when the true model was HCP). In the majority of simulations (87 out of 100, an example is shown in Supplementary Fig. 6E), both FCC and HCP models showed positive grid scores. Unlike the high SNR case where a true model showed higher grid scores and R square, grid scores were comparable for the two models in 76 cases and only 11 cases showed a higher grid score for the correct FCC model. R-squares were also similar in most cases. This means that the two models could not be distinguished. Of note, there were 8 cases where only the FCC model was significantly positive, and the grid score of the HCP model was not (Supplementary Fig. 6F). These simulation cases resonated with the actual fMRI data we observed (only the FCC model was significant in the left entorhinal cortex). In 2 cases, only the wrong model (HCP) was significant.

### Discussion

Based on the simulation results, we predict that if the true grid signal is large, FCC and HCP models would be correctly identified using our numerical estimation methods even when the sampled direction are limited to 120 degrees. When the SNR is low, our analysis will pick up evidence of a 3D grid signal, but it will not permit identification of which model was the correct one.

---

**Supplementary Fig. 6.**
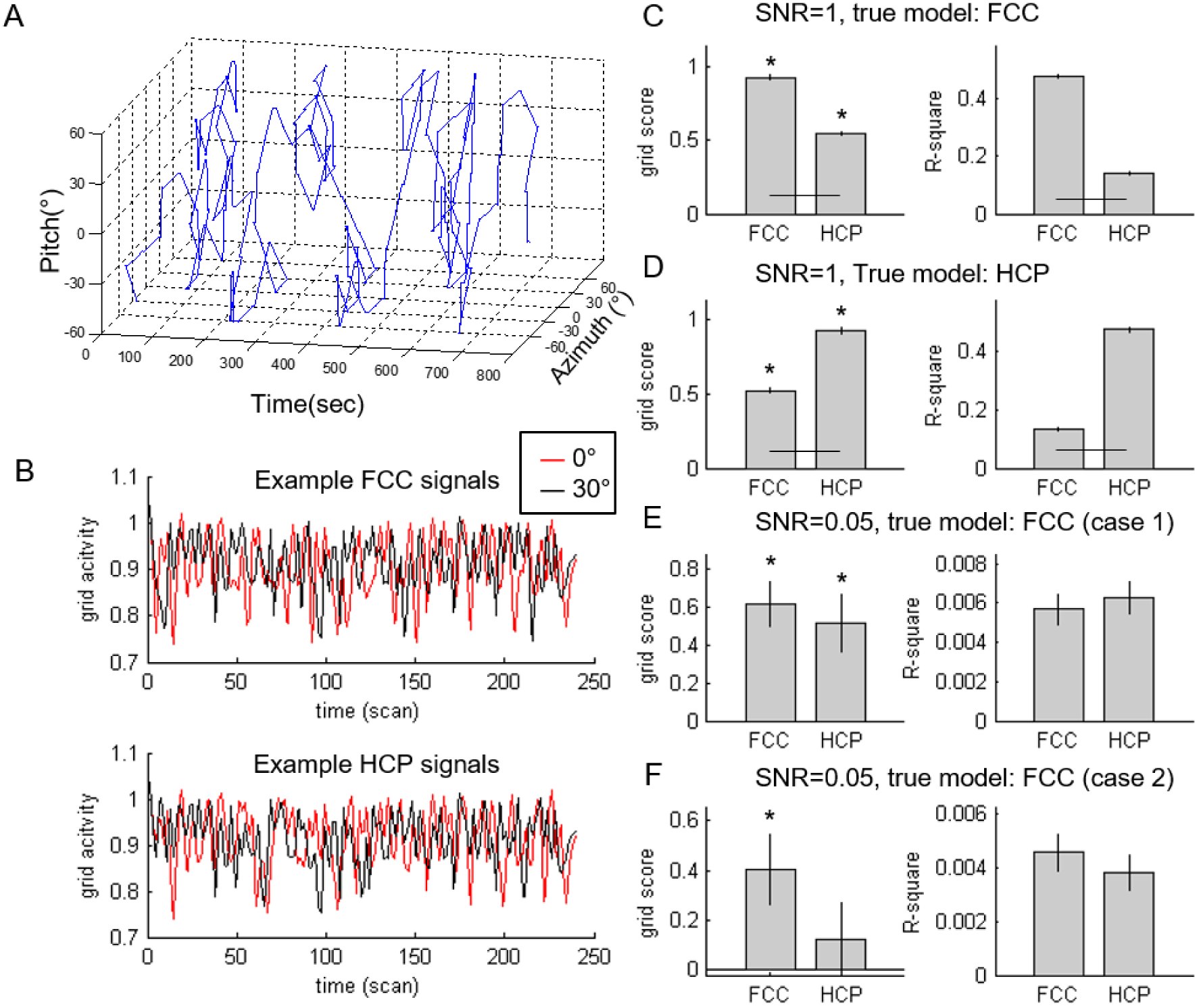
Simulation of the FCC and HCP models. **A.** A virtual 3D trajectory. Movement directions were limited to −60° to 60° vertically (pitch) and horizontally (azimuth). **B. A** hypothetical grid voxel responses when grid cells followed either the FCC (the top panel) or the HCP (the bottom panel) model. A grid cell’s activity is dependent on the grid orientation and movement direction. Two examples grid signals are shown when the orientation was 0° (red) or 30° (black). **C-F.** Simulation results when different SNR and different grid models were used. Thirty virtual participants were used. When the SNR was high, the FCC grid signal was correctly identified as the FCC model (higher grid score and R square for the FCC model compared to the HCP model) (**C.**), and the HCP grid signal was correctly identified as HCP model (higher grid score and R square for the HCP model) (**D.**). When the SNR was low, grid scores and R square were not significantly different for the FCC and HCP model fits (**E,F.**). In some cases, only the correct model showed a significantly positive grid score (**F.**). *p<0.05.

